# Force-modulated structural landscape of the catch bonding F-actin crosslinker α-actinin-4

**DOI:** 10.64898/2026.03.04.709699

**Authors:** Alfred C. Chin, Fatemah Mukadum, Matthew J. Reynolds, Glen M. Hocky, Gregory M. Alushin

## Abstract

Catch bonds, noncovalent supramolecular interactions whose lifetimes are increased by force, are ubiquitous in mechanical signaling pathways. The structural mechanisms of catch-bonding proteins remain unclear, hampering efforts to decipher how they are dysregulated in disease and exploit them therapeutically. The crosslinker α-actinin-4 (ACTN4) forms catch bonds with actin filaments (F-actin) to support the function of kidney podocytes, and its force-insensitive K255E variant causes autosomal dominant focal segmental glomerulosclerosis (FSGS). Using cryo-electron microscopy (cryo-EM), we find that wild-type ACTN4 engages F-actin in two modes, which biochemical experiments and molecular dynamics simulations assign as strong- and weak-binding states, while K255E ACTN4 only populates the strong binding state. By implementing a cryo-EM platform for applying tension across crosslinker–F-actin interfaces using myosin motors, we find that force promotes a weak-to-strong binding transition for wild-type ACTN4, consistent with a two-state catch bond model. Beyond providing mechanistic insight into how the K255E mutation disrupts ACTN4 F-actin catch-bonding in FSGS, this approach enables structural dissection of force-sensitive actin-binding proteins.

## Introduction

Catch bonds, noncovalent interactions which are counterintuitively strengthened by mechanical force^1^, are ubiquitous across mechanical signal transduction (mechanotransduction) pathways mediating cell adhesion^2–6^, cytoskeletal dynamics^7–9^, and immunity^10,11^. Malfunction of catch bonding proteins underlies hereditary diseases including focal segmental glomerulosclerosis (FSGS)^12,13^, while catch bond engineering can potentially optimize cell-based immunotherapies^14^. Structural mechanisms of catch bonds have not been directly visualized; instead, they have been extrapolated from theoretical models^1,15,16^, simulations^17–19^, and structural studies in the absence of force^20–22^. This has limited mechanistic insights into diseases associated with dysfunctional mechanotransduction and the corresponding development of therapeutics targeting mechanical signaling pathways.

Several actin-binding proteins (ABPs) form catch bonds with actin filaments (F-actin) to coordinate mechanotransduction through cell adhesion complexes and modulate biophysical properties of cells such as polarity and elasticity^2–4,7,23^. Optical trapping studies of how force impacts F-actin binding by the adhesion ABPs α-catenin, vinculin, and talin, all of which feature structurally homologous actin-binding domains (ABDs)^21,22,24,25^, are consistent with a two-state catch bond model^2–4^. This model features a short-lived weakly bound state and a long-lived strongly bound state^1^, with force promoting a transition from the weak state to the strong state (which is disfavored in the absence of force) to stabilize the interaction. While this model implies the existence of two F-actin-binding modes for these proteins, weak states have never been observed, with existing structures thought to represent biochemically-stabilized strong states^21,22,24,25^.

Moreover, force-dependent transitions between F-actin–bound states have also not been visualized, as determining structures of protein complexes in the presence of mechanical force has been infeasible. The structural underpinnings of the two-state catch bond model thus remain unclear, and it is unknown whether it generalizes to additional classes of catch-bonding ABPs beyond adhesion proteins.

The F-actin crosslinker α-actinin-4 (ACTN4) forms catch bonds with F-actin under physiological force regimes^13^ to support ACTN4’s functions in cell mechanics^7–9^. ACTN4 mutations cause podocytopathies, including the widely studied K255E variant that drives autosomal dominant, steroid-resistant FSGS^26^. K255E disrupts ACTN4 catch bonding by inducing constitutive high-affinity F-actin binding, resulting in binding lifetimes that monotonically decrease with force^7–9,13^ (“slip bonding”, a behavior that is typical for most protein-protein interactions^1^). Expression of K255E ACTN4 produces disorganized F-actin networks that disrupt the mechanical properties of podocytes, which are exposed to cyclic hemodynamic stresses during blood filtration by the kidney^12,27,28^. Despite extensive scrutiny, the pathophysiology of ACTN4 mutations remains unclear, limiting treatment options for ACTN4-associated FSGS.

Functioning as a rod-like homodimer mediated by spectrin repeats (Fig. 1A), ACTN4 crosslinks F-actin through its two tandem calponin-homology domains (CHDs; CH1 and CH2), a prevalent class of ABD in spectrin superfamily proteins^29^. Biochemical, computational, and structural studies have produced conflicting models for how α-actinins and other tandem CHD proteins engage F-actin, as previous negative stain and cryo-electron microscopy (cryo-EM) studies featured ambiguities in their interpretation of low resolution density maps^30,31^. Crystal structures of isolated tandem CHDs display a compact architecture, with a substantial interface between CH1 and CH2^32–35^. More recent high-resolution cryo-EM reconstructions have consistently shown that CH1 contacts F-actin, while CH2 is distal from the filament and is only weakly resolved, suggesting its orientation relative to CH1 becomes flexible upon F-actin engagement^36,37^. K255E and analogous high-affinity mutations in tandem CHD proteins^33,34,37,38^ are localized at the interface between CHDs, positioning them to impact inter-CHD dynamics. Collectively, these data support a model in which tandem CHDs must “open” to engage F-actin, which can be promoted by mutations that weaken CH2’s negative allosteric regulation of F-actin binding through CH1.

**Figure 1:**
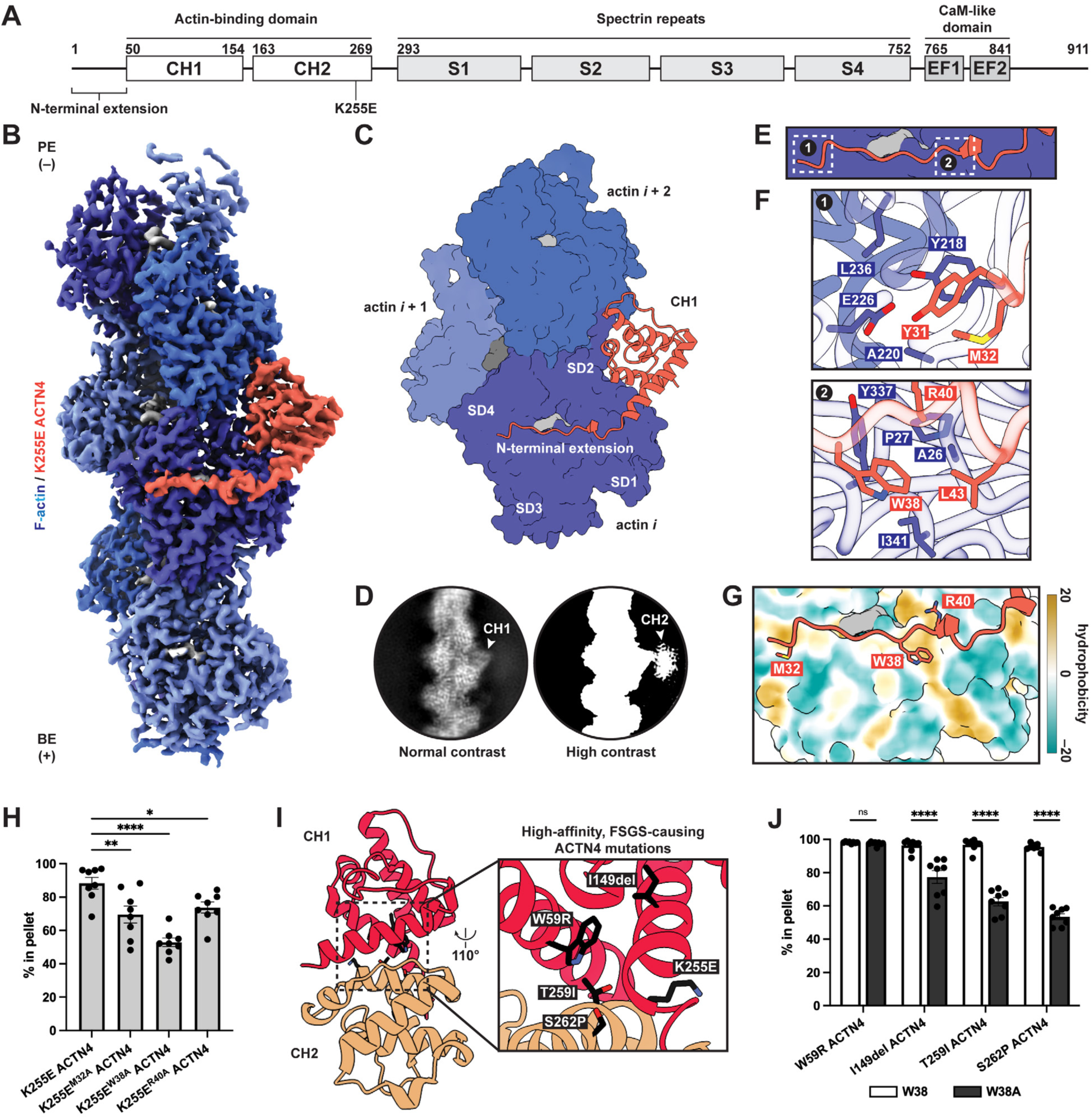
Structural basis for constitutive strong F-actin binding by K255E ACTN4. **(A)** Domain architecture of human ACTN4. **(B)** Cryo-EM map of K255E ACTN4 bound to F-actin. Phalloidin (dark gray) and ADP-P_i_ (light gray) are displayed. BE (+), barbed (plus) end; PE (−), pointed (minus) end. **(C)** Atomic model of K255E ACTN4 (cartoon representation) bound to F-actin (surface representation). SD, subdomain. **(D)** 2D class average of particles contributing to the cryo-EM reconstruction resolves diffuse CH2 density at low threshold. **(E)** Overview of interface between ACTN4’s NTE and actin. **(F)** Top: Detailed view of interactions at the extreme N-terminus (Y31 / M32) of the ordered portion of the NTE. Bottom: Detailed view of K255E ACTN4 W38 engaging a hydrophobic patch on actin’s surface. **(G)** View of actin-interacting residues in K255E ACTN4’s NTE. Actin is displayed in hydrophobic surface representation. Tan color indicates greater hydrophobicity. **(H)** Quantification of F-actin co-sedimentation assay for alanine mutations in the K255E ACTN4 NTE (representative gel in Fig. S4I). Conditions were compared by ordinary one-way ANOVA with Dunnett’s correction (*n* = 8). **(I)** Crystal structure of wild-type ACTN4 ABD^35^ (PDB 6O31) displaying high-affinity mutations that cause FSGS. **(J)** Quantification of F-actin co-sedimentation assay for high-affinity, FSGS-causing ACTN4 mutations W59R, I149del, T259I, and S262P and their corresponding W38A versions (representative gels in Fig. S4K). Conditions were compared by ordinary two-way ANOVA with Šídák’s correction (*n* = 8 per condition). For all co-sedimentation assay quantifications throughout the figure, data are presented as mean ± SEM. Individual data points are also displayed. \**P* < 0.05, \*\**P* < 0.01, \*\*\*\**P* < 0.0001, ns = not significant (*P* ≥ 0.05).

Mechanistic gaps nevertheless remain in our understanding of CHDs. Inter-CHD mutations alone are insufficient to drive opening of tandem CHDs in the absence of F-actin^33,34,37^, suggesting additional regulatory mechanisms are required to relieve auto-inhibition. Furthermore, previous studies have generally employed high-affinity mutants to achieve high-resolution F-actin bound structures, which have solely been obtained for isolated ABD constructs under saturation-binding conditions^36,37,39^. It thus remains unclear how tandem CHDs engage F-actin in the context of full-length wild-type proteins, and it is unknown whether and how ACTN4’s F-actin interface is modulated by force to mediate catch bonding.

## Results

### The K255E ACTN4–F-actin interface

To elucidate the F-actin engagement and force-regulatory mechanisms of ACTN4, we pursued cryo-EM structural studies. In this work, we employ recombinantly expressed full-length human ACTN4 proteins under sparse binding conditions mimicking those found in F-actin networks (Fig. 1A; Fig. S1A,B). We adapted our previously reported neural network-based particle picker for analyzing F-actin crosslinkers^40,41^ to localize interfaces between the rod-like protein and F-actin in cryo-EM micrographs (Fig. S2A,B; Methods).

We first examined the K255E variant, hypothesizing that its mechanosensing deficiency could result from the protein constitutively adopting a strong F-actin binding state. We determined a 3.2 Å resolution cryo-EM structure of the K255E ACTN4–F-actin interface using an asymmetric single-particle approach (Fig. 1B,C; Fig. S3A-D; Methods). This yielded a reconstruction of a single ABD bound to otherwise undecorated F-actin, consistent with the low abundance of K255E ACTN4 in our sample. Similar to previous cryo-EM structures of high-affinity tandem CHD mutants under saturation binding conditions, this map features well-defined density for CH1 contacting F-actin, while 2D class averages of particles contributing to the map display diffuse CH2 signal (Fig. 1D). From a pilot dataset using a higher concentration of K255E ACTN4, we also obtained a low-resolution cryo-EM map featuring weak CH2 density consistent with an open conformation (Fig. S4D-F). This resembles an F-actin-bound reconstruction of the high-affinity filamin A variant E254K, where CH2 opening resolves a steric clash with F-actin^36^. Consistently, the closed conformation observed in the K255E ACTN4 ABD crystal structure is incompatible with our cryo-EM reconstruction, as CH2 clashes with F-actin (Fig. S4G). Although an AlphaFold3 model recapitulates some features of the actin-binding interface, it does not predict CH2 opening (Fig. S4H).

CH1 displays only modest conformational changes upon F-actin binding versus the crystal structure of the isolated K255E ACTN4 ABD^34^, primarily in the N-terminal helix (Fig. S3E,F), indicating CH2 acts as a steric blocker of CH1 in the closed conformation rather than allosterically modulating CH1’s actin-binding surface. To validate the K255E ACTN4 F-actin binding interface, we performed F-actin co-sedimentation assays in the presence of the Lifeact peptide, which engages a site overlapping with CH1^42^ (Fig. S4A). Lifeact reduced F-actin binding by K255E ACTN4, consistent with competitive binding (Fig. S4B).

F-actin features inherent structural plasticity^43^, primarily focused at the D-loop of actin subdomain 2^44,45^, which mediates interactions between longitudinally adjacent subunits. We previously observed that the D-loop can adopt two primary conformations in bare F-actin, “in” and an “out”, which differ in the positioning of residues 45-50^44^. Previous structures of E254K filamin A^36^ and utrophin^39^ bound to F-actin show the D-loop to be stabilized in the “out” conformation (Fig. S3G). However, we find the D-loop to be stabilized in the “in” conformation by K255E ACTN4 (Fig. S3G,H). This suggests that, despite the similarity of their binding poses, different CHD ABDs can exert subtly different impacts on F-actin’s inherent structural landscape. This may also be linked to the different F-actin nucleotide states and stabilizing drugs used across studies^46^.

Our cryo-EM structure additionally resolved a portion of the ACTN4 N-terminal extension (NTE), which is disordered in solution and had been truncated to solve ACTN4 ABD crystal structures^34,35^. This segment forms an extensive interface with actin, extending across the nucleotide cleft to subdomain 4 (Fig. 1C). The first NTE residue that becomes ordered is Y31, phosphorylation of which inhibits F-actin binding^47^ (Fig. 1E). While other tandem CHD ABPs also feature NTEs that become ordered upon F-actin binding^36,37,39^, the extent of their engagement is limited compared to K255E ACTN4 (Fig. S4C). To assess the NTE’s contribution to F-actin binding, we mutated well-resolved actin contacting residues (M32, W38, and R40) and performed F-actin co-sedimentation assays. While mutating any of these residues to alanine significantly impaired K255E ACTN4 F-actin binding, W38A displayed the most substantial reduction (Fig. 1F-H; Fig. S4I,J). W38 is highly conserved and buried in a hydrophobic pocket on the actin surface, consistent with it playing a major role in F-actin binding.

We next examined other high-affinity ACTN4 mutants that cause FSGS^26,38^ (Fig. 1I), reasoning that if they employ an equivalent actin-binding interface, W38 would also be important for their F-actin engagement. Introducing the W38A mutation similarly inhibited F-actin binding in I149del, T259I, and S262P ACTN4, suggesting these FSGS mutations likely promote high-affinity F-actin binding through a similar mechanism as K255E. (Fig. 1J; Fig. S4K). However, W59R ACTN4 was unaffected by the W38A substitution, suggesting this variant may adopt a distinct F-actin binding mode. Collectively, these data show full-length K255E ACTN4 displays a similar F-actin-binding interface as other tandem CHD ABPs, featuring a key NTE-mediated actin contact that is shared with other high-affinity ACTN4 disease mutants^36,37^.

### Wild-type ACTN4 binds F-actin in weak and strong states

We next examined wild-type ACTN4, hypothesizing the protein might exhibit additional binding modes, such as a weak-binding state as anticipated by the two-state catch bond model. In our cryo-EM micrographs, wild-type ACTN4 formed thick, colinear F-actin bundles while K255E ACTN4 formed thin, disordered F-actin bundles (Fig. S5A), corroborating previous observations from negative stain electron microscopy^12^. Previous attempts to determine cryo-EM structures of wild-type isolated ABD constructs from β-III-spectrin and filamin A were hindered by poor F-actin decoration^36,37^, while helical processing of the wild-type ACTN2 ABD bound to F-actin under saturation binding conditions yielded a 16 Å cryo-EM map with density consistent for a single CHD^31^. Enabled by advances in automated data collection and our particle-picking approach (Fig. S2A,B), we successfully obtained a sufficiently large wild-type full-length ACTN4 dataset for single-particle analysis despite the protein’s low F-actin binding affinity and corresponding sparse distribution in micrographs.

From the same dataset, 3D classification resolved two primary binding modes (Fig. 2A; Fig. S5B). Class one, which we refined to 4.1 Å resolution, features wild-type ACTN4 in a conformation indistinguishable from that we observed for K255E ACTN4, with ABD density wedged in the cleft between longitudinally adjacent actin subunits (Fig. 2A, left; Fig. S6A-E). Class two, refinement of which stalled at ∼12 Å resolution for the ABD region of the map (Fig. 2A, right; Fig. S6F-H), was entirely distinct. This class features a larger ABD density, consistent with the size of both CHDs, contacting the N-terminal regions of adjacent actin subunits along the same protofilament (Fig. 2A, right; Fig. S6F-J). As the well-resolved class one structurally resembled K255E, we tentatively assigned this as a strong-binding state, whereas the poor resolvability of class two is suggestive of a flexible, weakly bound state.

**Figure 2:**
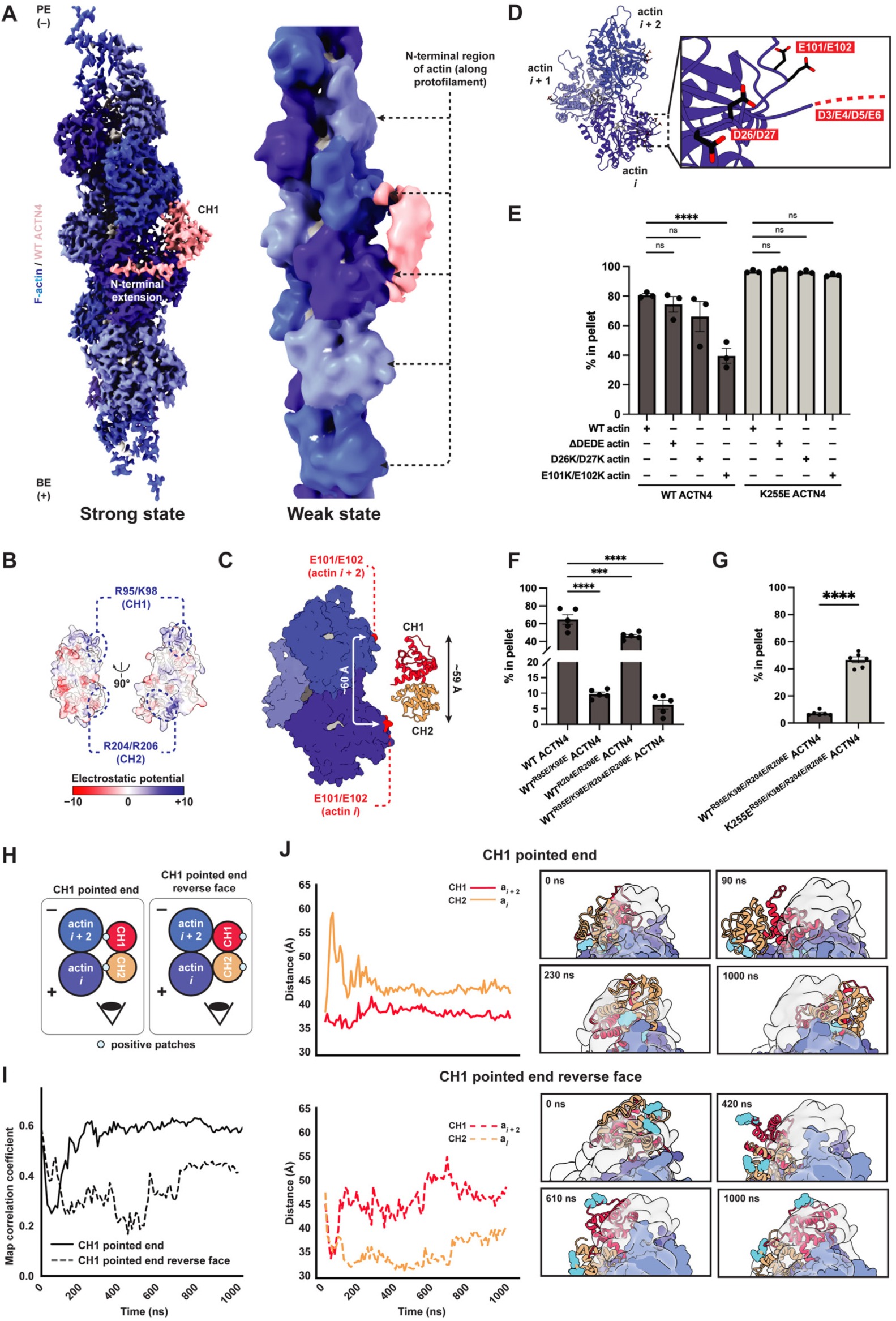
Two-state F-actin binding by wild-type ACTN4. **(A)** Cryo-EM maps of wild-type ACTN4 bound to F-actin in strong (left) and weak (right) states. Weak state map is low-pass filtered to 12 Å. Phalloidin (dark gray) and ADP-P_i_ (light gray) are displayed. BE (+), barbed (plus) end; PE (−), pointed (minus) end. **(B)** Detailed view of negative patches adjacent to the N-terminus of each actin subunit in F-actin^89^ (PDB 9DUU). D3-E6 are flexible and unresolved in most actin structures. **(C)** Quantification of F-actin co-sedimentation assay for wild-type (WT) and K255E ACTN4 in the presence and absence of charge reversal mutations in recombinant human α-actin-1 (representative gels in Fig. S7E). Conditions were compared by ordinary two-way ANOVA with Dunnett’s correction (*n* = 3). **(D)** Distance between E101/E102 of neighboring human α-actin-1 subunits along the same strand compared to the span of the long axis of the wild-type ACTN4 ABD^35^ (PDB 6O31), measured as the distance between G102 and K181. **(E)** Electrostatic potential surface map of wild-type ACTN4 ABD. **(F)** Quantification of F-actin co-sedimentation assays for charge reversal mutations in the context of wild-type ACTN4 (representative gel in Fig. S7F). Conditions were compared by ordinary one-way ANOVA with Dunnett’s correction (*n* = 5). **(G)** Quantification of F-actin co-sedimentation assay for combined charge reversal mutations in both CHDs of wild-type and K255E ACTN4 (representative gel in Fig. S7G). Conditions were compared by unpaired t test (*n* = 6). (**H**) Cartoon of ACTN4 ABD orientations examined in molecular dynamics (MD) simulations of weak state. **(I)** Map correlation coefficients from 1 µs MD simulations of ACTN4 ABD orientations from **(H)**. **(J)** Plots of CH domain distance to the nearest actin subunit (left) and representative simulation snapshots overlaid with the cryo-EM map (right) of the CH1 pointed end (top) and CH1 pointed end reverse face (bottom) orientations. Snapshots are from the view indicated in **H**. Distances were calculated between the centers of mass of C_α_ atoms. Cyan residues indicate positive amino acids highlighted in **B**. For all co-sedimentation assay quantifications throughout the figure, data are presented as mean ± SEM. Individual data points are also displayed. \*\*\**P* < 0.001, \*\*\*\**P* < 0.0001, ns = not significant (*P* ≥ 0.05).

To probe the binding modes of wild-type ACTN4, we first introduced perturbations that inhibited F-actin binding by K255E ACTN4, anticipating these would impact the putative strong-binding state. Consistently, competition with Lifeact and the W38A mutation both also impaired F-actin binding by wild-type ACTN4 (Fig. S7A,B). Deleting the entire NTE also reduced F-actin binding, albeit to a lesser degree than the W38A mutation (Fig. S7B), an observation consistent with previous reports of autoinhibitory elements in this segment^47,48^. Completely removing the NTE in the presence of the K255E mutation produced an unstable construct which aggregated; thus, we were unable to assess the interplay of these two lesions.

We next sought to identify perturbations that selectively inhibit F-actin binding by wild-type ACTN4, reasoning they would impact the putative weak state. Consistent with a role for actin’s N-terminal region, treating actin with subtilisin^49^, a protease which cleaves between amino acids 47 and 48 of the D-loop, inhibited F-actin binding by wild-type but not K255E ACTN4 (Fig. S7C,D). This observation additionally suggests ACTN4’s stabilization of the “in” D loop conformation (Fig. S3G,H) does not play an important role in the formation of the strong F-actin binding state. Since the precise impact of subtilisin cleavage on F-actin structure is unknown, we next sought to identify specific contacts selectively mediating binding by wild-type ACTN4. Examining the structure of ACTN4’s ABD in the closed conformation revealed a positive patch at the tip of each CHD, aligned along one surface of the molecule (Fig. 2B). The overall length of the ABD (∼59 Å) closely matches the distance between the subdomain 1s of longitudinally adjacent actin subunits, which feature a negative patch (including the negatively charged actin N terminus) previously implicated in mediating contacts with myosin during the weak-binding phase of the motor’s mechanochemical cycle (Fig. 2C,D)^50,51^. We thus hypothesized that weak F-actin binding by wild-type ACTN4 could be mediated by electrostatic interactions between these complementary charged patches.

To test this hypothesis, we mutated both surfaces and performed co-sedimentation assays. We first prepared recombinant actin featuring an N-terminal truncation (ΔD3/E4/D5/E6, which we refer to as ΔDEDE) or charge reversal mutations (D26K/D27K and E101K/E102K, Fig. 2D). While ΔDEDE and D26K/D27K had no clear effect, the E101K/E102K double mutation significantly reduced F-actin binding by wild-type ACTN4, while none of the mutations affected F-actin binding by K255E ACTN4 (Fig. 2E; Fig. S7E). This suggests actin residues E101/E102 specifically mediate ACTN4’s weak F-actin binding.

We next introduced charge reversal mutations into the positive patches of ACTN4 (R95E/K98E in CH1 and R204E/R206E in CH2). Both lesions significantly reduced F-actin binding, with the R95E/K98E CH1 mutations displaying a substantially more potent effect, suggesting the CH1 patch plays a more prominent role in F-actin engagement (Fig. 2F; Fig. S7F). The quadruple mutant (R95E/K98E/R204E/R206E) nearly eliminated F-actin binding (Fig. 2F; Fig. S7G). Binding was significantly rescued by introducing the K255E mutation in the quadruple charge reversal background (R95E/K98E/R204E/R206E/K255E), suggesting a selective effect primarily on wild-type ACTN4’s weak F-actin binding state (Fig. 2G; Fig. S7G).The CH2 mutations R204E/R206E in isolation had no detectable effect on F-actin binding by K255E ACTN4 (Fig. S7H), consistent with CH2 being displaced from the filament surface in our K255E ACTN4–F-actin reconstruction (Fig. 1D; Fig. S4D-F), while the CH1 mutations R95E/K98E moderately reduced F-actin affinity (Fig. S7H). A potential trivial explanation for this observation would be R95/K98 participating in the CH1-mediated strong state F-actin binding interface, but these residues are instead oriented away from F-actin in our K255E ACTN4–F-actin reconstruction (Fig. S7I). Future studies will thus be required to ascertain how the ACTN4 CH1 positive patch impacts K255E ACTN4’s F-actin engagement. Our data are nevertheless broadly consistent with the presence of a weak F-actin binding mode in wild-type ACTN4, mediated by electrostatic interactions between positive patches in both CHDs and negatively charged patches on the F-actin surface.

### Molecular dynamics simulations of the weak state

Due to the low resolution of our weak state cryo-EM reconstruction, we pursued molecular dynamics (MD) simulations to probe the interaction between F-actin and the tandem CHDs in greater detail. Guided by our density map, we manually placed the crystal structure of the wild-type ACTN4 ABD^35^ (PDB 6O31), which is effectively indistinguishable from the K255E ACTN4 ABD crystal structure (Fig. S8A,B), adjacent to the F-actin surface, with the ABD positive patches approximately positioned in proximity to the negative actin patches. As a negative control, we also rotated the ABD 180° about its long axis, such that the positive patches were facing away from F-actin (“reverse face”). As our density map features insufficient resolution to discriminate the individual CHDs, we prepared models with both CH1 (CH1 PE) and CH2 (CH2 PE) oriented towards the pointed (minus) end of F-actin (Methods). We then performed all-atom MD simulations with explicit solvent following established protocols for ABP–F-actin complexes^19,52^ (Fig. 2H-J; Fig. S8C-F; Methods), without using the cryo-EM map as restraint. After performing a microsecond simulation for each initial pose to allow the system to sample possible stable configurations, we performed four 500 ns replicate simulations for both CH1 PE and CH2 PE, with starting structures for the replicates taken from equally spaced intervals along the corresponding initial simulation trajectory (Fig. S8C).

In the CH1 PE orientation, the ABD formed contacts with F-actin through both CHDs, while in the reverse face orientation it tumbled randomly without clearly engaging the filament (Fig. 2H,J; Movie S1). We then analyzed the dynamics of the CH1 PE complex by calculating the distance between each CHD and its contacting F-actin over simulation frames for each replicate (Fig. S8C, top; Fig. 2J). We observe that CH1 remains consistently in contact with F-actin (Movie S1), indicated by a stable short distance, while CH2 repeatedly binds and unbinds the F-actin surface, indicated by oscillations in the distance trace (Fig. S8C, top; Fig. 2J). This is consistent with our biochemical data suggesting that positive residues in CH1 make a greater contribution to F-actin binding by wild-type ACTN4 than those in CH2 (Fig. 2F). To assess the consistency between simulations and our experimental density map, we extended the CH1 PE and CH2 PE replicates featuring the smallest fluctuations in CHD distances to actin (marked by asterisks in Fig. S8C,D), indicating stable poses, for an additional 500 ns. We then quantified the overlap between ABD atoms and the cryo-EM map. Across simulation frames, the CH1 PE complex showed consistently higher correlation with the map than the reverse face (Fig. 2I), providing additional support for this binding orientation.

The CH2 PE complex (Fig. S8D) displayed similar dynamics in two out of four replicates, with CH1 forming a stable contact with F-actin and CH2 engaging in only transient interactions (Fig. S8C, bottom; Fig. S8F, top). However, in the remaining two replicates, both CHDs remained in consistent close contact with F-actin, in an orientation that is inconsistent with our mutagenesis data. We observed a similar phenomenon for the CH2 PE reverse face (Fig. S8F, bottom) which may represent either an uncharacterized metastable minimum that is sparsely populated under the experimental conditions we employed or a simulation artifact (Movie S1). We furthermore observed an even higher stable map correlation for the CH2 PE reverse face versus the CH2 PE complex (Fig. S8E), once again contradicting our biochemical data, while neither CH2 PE nor CH2 PE reverse face reached as high of a correlation plateau as the CH1 PE complex (Fig. 2I). Collectively, these simulation results support an ACTN4 weak F-actin binding mode in a closed ABD conformation mediated by electrostatic interactions through both ABDs, with the CH1 PE orientation displaying dynamics consistent with our structural and biochemical data.

### Reconstituting ACTN4–F-actin in the presence of motor forces

To assess how ACTN4’s F-actin–bound structural landscape is linked to catch bond formation, we next sought to determine how force impacts the ACTN4–F-actin interface. We adapted our recently reported approach for reconstituting myosin motor activity on holey carbon cryo-EM substrates^53^ to apply force across ACTN4-crosslinked F-actin filaments, placing the ACTN4-F-actin complex in a load-bearing geometry compatible with catch bond formation (Fig. 3A). We employed the plus-end directed motor myosin Va, as individual myosin V heads exhibit a stall force of ∼3 pN, such that small ensembles of dimeric motors on the grid surface can collectively apply sufficient load to activate ACTN4 catch bonding (a critical force of 4 pN)^13,54,55^.

**Figure 3:**
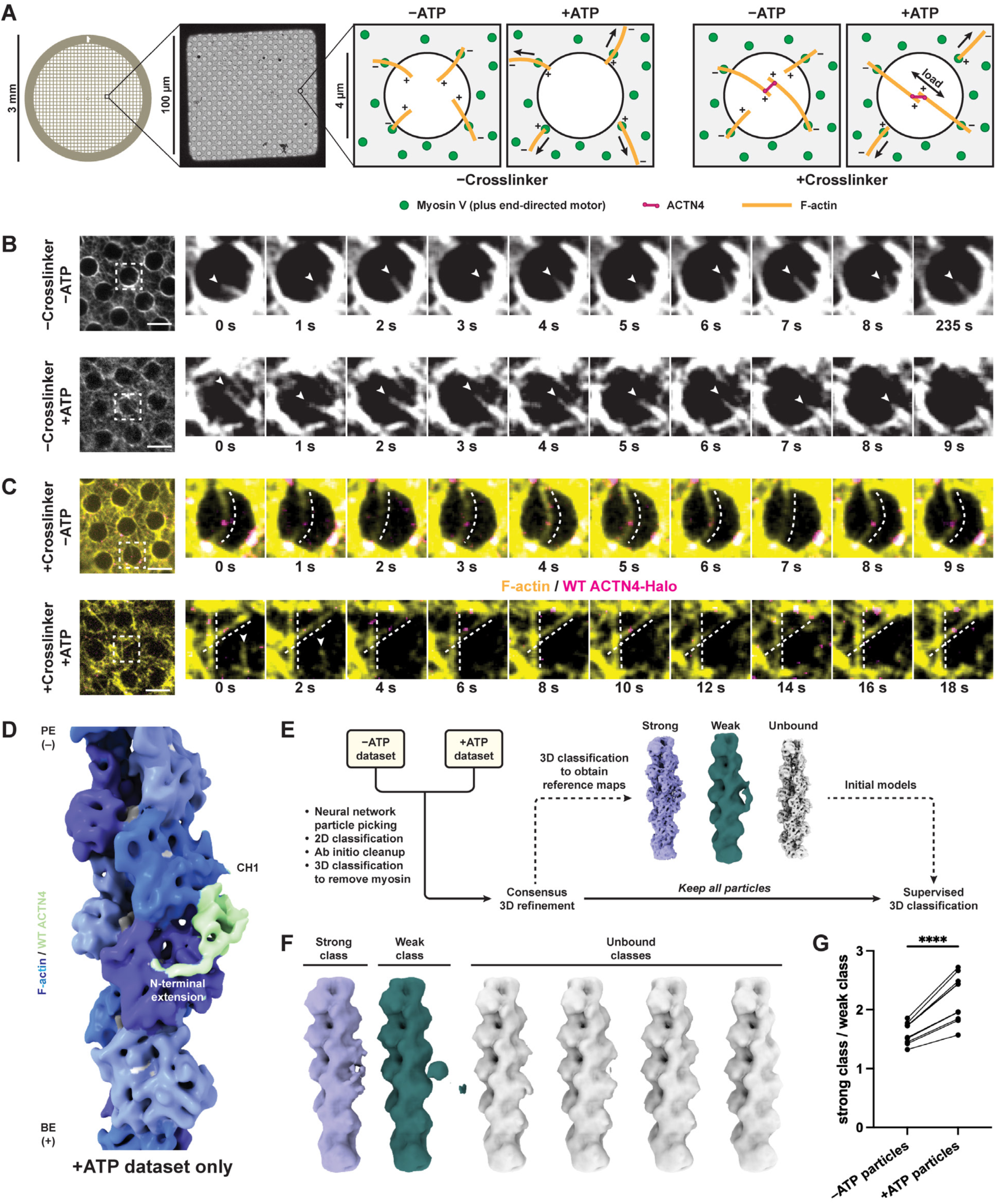
Myosin forces promote an ACTN4 weak-to-strong F-actin binding transition. **(A)** Schematic of myosin motor-based force reconstitution assay on cryo-EM substrates. **(B)** Epifluorescence of force reconstitution assay in absence of ACTN4 without (top) and with (bottom) ATP. F-actin (Alexa Fluor 568 phalloidin) channel is displayed. Arrowheads indicate filaments undergoing thermal fluctuations without ATP (top) and directed motility with ATP (bottom). Scale bars, 5 μm. **(C)** Epifluorescence of force reconstitution assay in presence of ACTN4 without (top) and with (bottom) ATP. F-actin (Alexa Fluor 568 phalloidin) and wild-type ACTN4-Halo (Janelia Fluor 646) channels are displayed. Dotted lines indicate F-actin bundles which are curved (slack) without ATP (top) and straight (taut) with ATP (bottom). Arrowheads indicate non-crosslinked F-actin gliding onto the carbon film in the presence of ATP. Scale bars, 5 μm. **(D)** Cryo-EM map of wild-type ACTN4 bound to F-actin from the +ATP dataset, which is consistent with the strong state. Phalloidin (dark gray) and ADP-P_i_ (light gray) are displayed. BE (+), barbed (plus) end; PE (−), pointed (minus) end. **(E)** Workflow for combined processing of −ATP and +ATP datasets for quantification through 3D classification. See Fig. S10B for additional details. **(F)** Representative class assignments from a 3D classification run. **(G)** Quantification of strong versus weak class assignments for particles derived from −ATP and +ATP datasets, presented as each dataset’s strong / weak particle number ratio per 3D classification run. Conditions were compared by ratio paired t test (*n* = 10 independent 3D classification runs). \*\*\*\**P* < 0.0001.

We reasoned that in the absence of ACTN4 crosslinker, the substrate geometry would favor ATP-dependent F-actin gliding and enrichment on the surface of the carbon film, as motors anchored to the film surround empty holes. The presence of ACTN4, which primarily crosslinks F-actin in an antiparallel orientation^12,56^, results in the formation of bundles. In the presence of ATP, anti-parallel bundles exposed to balanced opposing forces through their constituent filaments should come under tension (Fig. 3A), remaining suspended over holes. This geometry recapitulates dilation stress in actin networks, which specifically drives mechanosensitive subcellular ACTN4 enrichment in a catch bond-dependent manner^7,8^. Conversely, individual F-actin filaments and rare parallel bundles would be translocated on to the carbon surface (Fig. S9A). This self-organizing behavior selectively preserves tensed bundles within holes for cryo-EM imaging, mitigating the inherent heterogeneity of force-generating actin-myosin networks. Preparing specimens in the presence and absence of ATP to activate motors furthermore enables direct visualization of how motor forces modulate the distribution of conformational states in the system.

To calibrate the assay, we performed time-lapse epifluorescence microscopy using fluorescently labeled F-actin (Alexa Fluor 586 phalloidin) and wild-type ACTN4 featuring a C- terminal HaloTag labeled with Janelia Fluor 646, first examining specimens in the absence of ACTN4. Without ATP, no directed motility was observed, with unbundled F-actin primarily adhering to the carbon film through strong-binding interactions with rigor myosin motors. A subset of filaments protruded into holes, which exhibited random fluctuations (Fig. 3B, top; Movie S2). With ATP, F-actin exhibited gliding dynamics, with filaments being drawn on the carbon film, where they displayed extensive motility (Fig. 3B, bottom; Movie S2). In the presence of crosslinker, we observed the formation of bundles suspended over holes featuring wild-type ACTN4-Halo puncta (Fig. 3C). Without ATP, these bundles were slack, featuring substantial, fluctuating curvature (Fig. 3C, top; Movie S3). Conversely, upon ATP addition, we observed taut, straight bundles, in addition to gliding individual filaments on the carbon film (Fig. 3C, bottom; Movie S3). Quantification indicates a significant decrease in bundle curvature in the presence of motor activity, supporting an increase in bundle tension (Fig. S9B). We also observed mechanical breakage of bundles over time, consistent with sufficient forces being generated to rupture ACTN4–F-actin bonds (Fig. S9C). Collectively, these data support our successful reconstitution of ACTN4 crosslinking actin filaments bearing physiological motor forces in a format suitable for cryo-EM structural studies.

### Mechanical force promotes a weak-to-strong state transition

To determine how myosin-generated forces impact wild-type ACTN4’s F-actin engagement, we plunge froze force reconstitution assay specimens in the absence of ATP that were additionally treated with apyrase to remove nucleotide (without force, “–ATP”) and in the presence of ATP (with force, “+ATP”). We used an untagged construct for wild-type ACTN4 instead of the HaloTagged construct employed for epifluorescence microscopy, but we continued to include fluorescently labelled F-actin (Alexa Fluor Plus 555 phalloidin). This allowed us to screen grids using cryo-fluorescence microscopy (Fig. S9D) to verify that F-actin was present in holes after the grid preparation process, as bundles rupture over time (Fig. S9C). To determine if the presence of force enriched any prominent states that we had not observed in the absence of motor activity, we initially processed the +ATP dataset using our single-particle workflow. This procedure produced a 6.7 Å resolution reconstruction which closely resembled the K255E ACTN4 structure / putative wild-type ACTN4 strong-binding state observed in the absence of motors (Fig. 3F; Fig. S10A). While this result does not rule out the presence of rare and / or flexible states that cannot be recovered through averaging, it suggests that force does not primarily regulate ACTN4 by populating a unique conformation.

We next examined whether force shifted the conformational landscape we observed in the absence of motors. To minimize bias while assessing the impacts of force on this distribution, we shuffled particles from the −ATP and +ATP datasets and processed them jointly (Fig. 3G; Fig. S10B), adapting an approach previously applied to studying structural transitions in ion channels under different biochemical conditions^57^. Following 2D classification and *ab initio* cleanup, we performed 3D classification to discard particles containing myosin V, as a subset of motors were not attached to the carbon film. Refining these particles produced a 4.0 Å structure of a motor domain bound to F-actin (Fig. S10B), confirming their molecular identity. All remaining particles were retained for downstream analysis to maximize statistical rigor. Using these remaining particles and our single-particle procedure, we obtained reconstructions of strong, weak, and unbound states that were then used as references for supervised 3D classification (Fig. 3G; Fig. S10B).

We then performed 10 independent runs of supervised 3D classification with 6 classes (1 strong reference, 1 weak reference, and 4 unbound references) lowpass filtered to 15 Å. Based on visual inspection, each run correspondingly output 1 strong class, 1 weak class, and 4 unbound classes, consistent with the references (Fig. 3H). To assess the robustness of the classification procedure, we analyzed how particles originally contributing to the strong, weak, and unbound reference maps were assigned across classes (Fig. S10C-E). Strong reference particles were enriched in the strong class, while weak and unbound reference particles were enriched in the weak and unbound classes, respectively. To assess the reproducibility of assignments across 3D classification runs, we calculated the proportion of shared particles between 3D classes across all 10 runs (Fig. S10F). Strong classes shared a high proportion of particles with other strong classes. This was also true when comparing weak classes with each other, albeit with slightly more variability. However, strong classes shared a negligible proportion of particles with weak classes.

To determine whether force redistributes the populations of bound states, we then examined the source dataset of the particles contributing to each class. Examining the ratio of particles in the strong class versus the weak class in each dataset (Fig. 3I), we find that particles derived from the +ATP dataset consistently populated the strong class over the weak class, compared to particles from the −ATP dataset. Moreover, the fraction of particles derived from the +ATP dataset was higher in the strong class than the weak class (Fig. S10G). Comparing the enrichment of +ATP particles in each of the 6 classes revealed that the strong class consistently ranked in the upper half, whereas the weak class always ranked at or near the bottom (Fig. S10H).

We next evaluated the behavior of individual particles between classification runs. We profiled particle class assignments across all 10 runs, quantifying how many times each particle was assigned to a given class (Fig. S10I). For the strong class, we observed a bimodal distribution with peaks at 1, representing particles with flexible class assignment that only contributed to the strong class once, and 10, representing particles assigned to the strong class every time. A similar distribution was observed for the weak class (Fig. S10J). If particles were randomly assigned to classes, such a bimodal distribution would not be anticipated. We next binned particles by how many times they were assigned to each class and recalculated the strong / weak class ratio (Fig. 3I; Fig. S10K). Particles that consistently populated their respective class displayed an even more prominent shift towards the strong class when comparing the –ATP and +ATP conditions, while this effect was diminished in particles with variable class assignments. This analysis suggests that the force-dependent shift towards the strong state we observe is primarily driven by particles whose state can be reproducibly assessed through 3D classification. Overall, this analysis is consistent with wild-type ACTN4 featuring two F-actin binding states, with force promoting a population-level transition from the weak-binding to the strong binding state. It also supports the single-CHD bound interaction mode as the strong binding state, and the doubly-CHD bound interaction mode as the weak-binding state.

## Discussion

In this study, we have directly visualized two F-actin binding modes by the catch bonding actin crosslinker ACTN4 and shown that mechanical force promotes a population-level shift from a weakly bound state to a strongly bound state (Fig. 4). These data indicate that ACTN4’s force-regulated F-actin binding landscape is compatible with a two-state catch bond model, as previously proposed for cell adhesion proteins^2–4^, suggesting this may be a more general mechanism by which ABPs can form catch bonds with F-actin. We also show that the FSGS-causing K255E mutation locks ACTN4 in the strongly-bound state in the absence of force, providing a structural explanation for this variant’s constitutive high-affinity F-actin binding. By demonstrating the feasibility of using our myosin force reconstitution assay and cryo-EM to visualize the force-regulated structural landscapes of ABPs and their disruption by pathological mutations, this work furthermore provides a platform for more broadly dissecting the structural mechanisms of cytoskeletal mechanotransduction and its dysfunction in disease.

**Figure 4:**
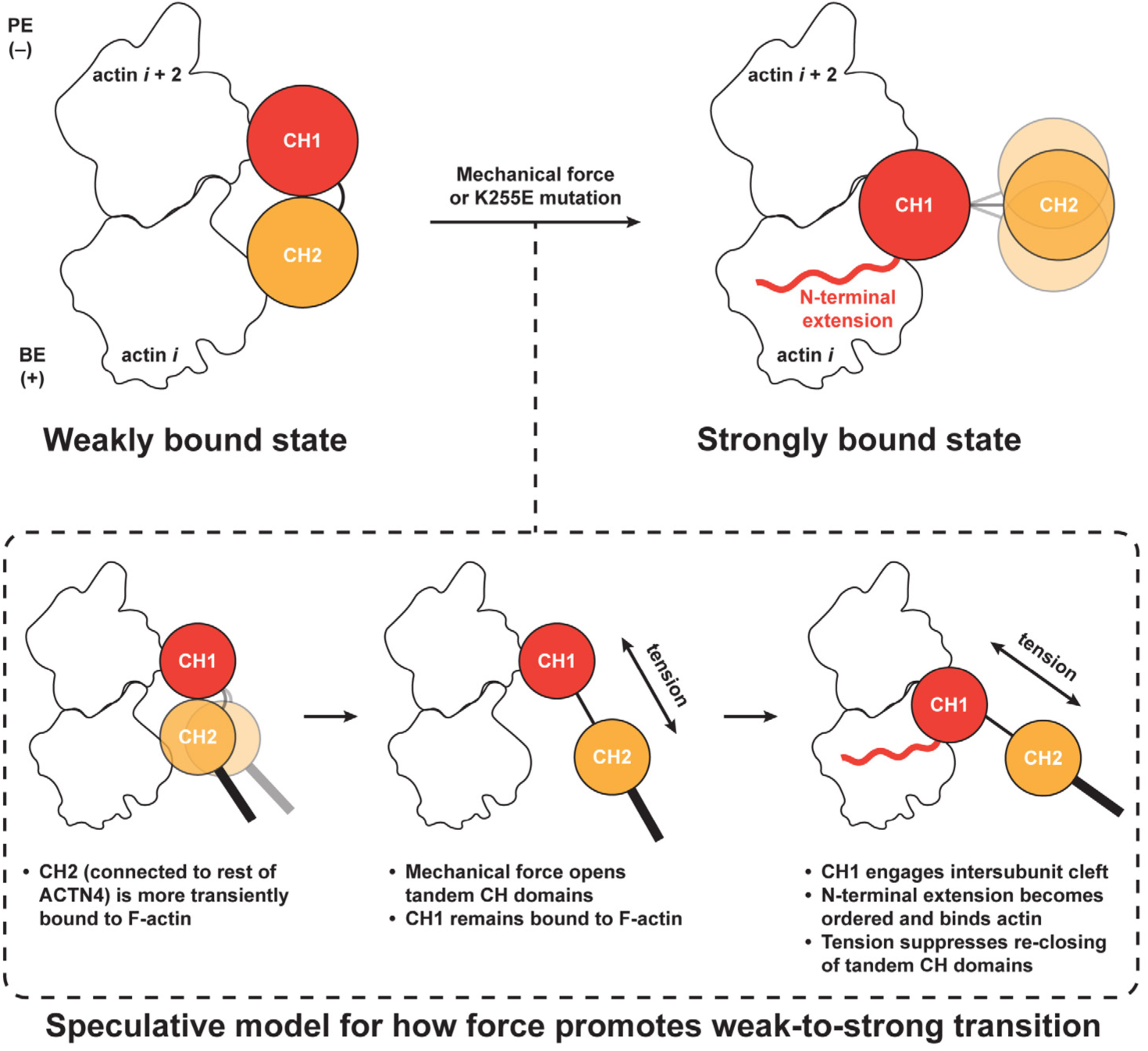
Proposed mechanism for force-regulated two-state F-actin binding by ACTN4. Wild-type ACTN4 binds F-actin in a weak state, with both CHDs engaging the N-terminal region of neighboring actin subunits on the same protofilament. In the presence of mechanical force or the K255E mutation, the tandem CHDs open, and the ABD adopts a strongly bound state in which CH1 docks into the cleft between actin subunits while the NTE undergoes a disorder-to-order transition to engage the filament surface. CH2 is unbound and highly flexible in the strongly bound state. While the detailed pathway underlying the force-mediated weak-to-strong binding transition remains to be determined, our charge reversal mutants and MD simulations indicate that CH2 engages F-actin more transiently than CH1 in the weak state. Because CH2 is connected to the rest of the ACTN4 molecule, we speculate that tension across the crosslinker can rupture the CH2**–**F-actin interaction. This would promote inter-CHD separation to open the ABD, freeing CH1 to bind F-actin in the strong state. Once the ABD is strongly bound, tension could also prevent CH2 from re-binding CH1 to close the ABD, further mediating stabilization of the strong state.

Leveraging advances in cryo-EM data processing for analyzing cytoskeletal crosslinkers^41,58^, we successfully determined structures of a wild-type tandem CHD ABD bound to F-actin in the context of its full-length protein. This uncovered a weak-binding state for ACTN4, mediated by simultaneous actin contacts through positive patches on both CHDs, which remain in a closed conformation (Fig. 2A; Fig. 4). Indirect biophysical studies and computational simulations have previously suggested that tandem CHDs could bind F-actin while closed, although these studies predicted different interfaces from those which we visualize here^56,59,60^. Our study reconciles this model with more recent cryo-EM work demonstrating CH2 separation and F-actin contact solely through CH1 by tandem CHDs^31,36,37^, which we delineate as the strong-binding state for ACTN4. Moreover, our findings provide an explanation for the previously counterintuitive finding that ABD truncation constructs featuring CH1 alone have a lower F-actin binding affinity than a full CH1-CH2 construct^31,61^, despite CH2 being auto-inhibitory in the context of ABD opening, since CH2 mediates F-actin contacts in the weak-binding state.

As tandem CHDs are widely distributed across the ABP proteome, we examined whether other ABDs featured an appropriate charge distribution to potentially adopt a weak-binding state through a similar binding pose. Positive patches on CH1 that mirror those on ACTN4 appear to be consistently conserved, but there is variation in the presence of positive patches on CH2 (Fig. S11A), suggesting this capacity may have diverged across the spectrin superfamily. Future studies will be required to examine whether a positive CH2 patch is correlated with F-actin catch-bonding activity in tandem CHD proteins, the generality of which remains unknown. In addition to its diversification across proteins, CH2 charge distribution may also be impacted by mutations. In filamin B, the high-affinity atelosteogenesis mutation W148R is localized in CH2, distal from the inter-CHD interface, and minimal conformational differences were observed between the wild-type and mutant ABDs^62^. We find that the W148R filamin B ABD features a CH2 positive patch that is absent from the wild-type (Fig. S11B), suggesting its increased F-actin binding affinity could be mediated by stabilization of a weak-binding state rather than enhanced ABD opening. We therefore propose that examining potential alterations to two-state F-actin binding is an important additional consideration when interpreting disease mutations in tandem CHD proteins.

Although our structural findings delineate the strong and weak F-actin binding states of ACTN4, they do not directly shed light on how force promotes a weak-to-strong transition. We observe that CH2 makes a relatively minor contribution to F-actin binding by wild-type ACTN4 (which we interpret to occur in the weak binding state; Fig. 2F), and CH2 repeatedly binds and unbinds in our simulations of the weak-binding state, while CH1 remains stably bound (Fig. 2J; Movie S1). This leads us to speculate that tension across the ACTN4 molecule can promote CH2 dissociation from F-actin and inter-CH opening while CH1 remains engaged (Fig. 4). This could facilitate ABD opening and favor formation of the strong-binding state, which could occur either through force-mediated sliding of CH1 along F-actin to the inter-actin subunit cleft or by CH1 unbinding and rebinding, favored by the high local concentration of the ABD, particularly if the other end of the crosslinker remains bound to a nearby filament.

A limitation of our weak-state MD simulations is our omission of the ACTN4 NTE (due to a lack of experimental structural information for this segment in free ACTN4 or in the weak-state density map), given that our structural and biochemical data show it plays a key role in the strong-binding state (Fig. 1; Fig. S7B). The NTE may also contribute to the weak-to-strong transition, and studying its role is an important topic for future work. Additionally, the ambiguity of the ABD orientation in the weak state may give rise to different force geometries, with the potential for multiple pathways for transitioning to the strong state. Nevertheless, the convergence between our structural data, biochemical data, and simulations favor the CH1 PE orientation. This pose is also consistent with a recent report that the cardiac α-actinin ACTN2 (an ACTN4 paralog that also features a CH2 positive patch; Fig. S11A) forms a directional catch bond with F-actin that is selectively stabilized by minus (pointed)-end directed force^63^. In our proposed scheme, the CH1 PE orientation is compatible with CHD opening by tension in this force geometry (Fig. 4), which is also anticipated to be the dominant geometry in our force reconstitution assays (Fig. 3A). Once the ABD has adopted the strong state, tension across the crosslinker would prevent CH2 from re-engaging CH1 to close the ABD and promote its dissociation from F-actin, providing an additional potential mechanism for force-dependent stabilization (Fig. 4). Future biophysical studies, as well as MD simulations incorporating force application, will be necessary to fully define the dynamics of ACTN4 F-actin catch-bonding, building upon the structural foundation we have provided here.

## Acknowledgements

This work was funded by a Rockefeller University Center for Clinical and Translational Science (NCATS CTSA #UL1TR001866) Pilot Grant to A.C.C. and NIH / NIGMS grant R35GM161251, a Rockefeller University Anderson Cancer Research Award, a Gabrielle H. Reem and Herbert J. Kayden Early-Career Innovation Award, and a Black Family Therapeutic Development Fund Pilot Award to G.M.A. Additionally, G.M.A. is a Biohub Investigator. A.C.C. was supported by NIH Medical Scientist Training Program Training Grant T32GM152349, an American Heart Association Predoctoral Fellowship, a Croucher Foundation Scholarship for Doctoral Study, and a NIDDK F30 Ruth L. Kirschstein National Research Service Award (F30DK138769). GMH was supported by NIH / NIGMS grant R35GM138312. FM was supported by an NSF GRFP award. Simulation work was supported in part through NYU Information Technology High Performance Computing resources, services, and staff expertise. We thank H. Ng, J. Sotiris, and M. Ebrahim for assistance with cryo-EM/cryo-fluorescence data collection at Rockefeller’s Evelyn Gruss Lipper Cryo-Electron Microscopy Resource Center, A. Pan and L. Sutton for assistance with actin purification, R. Gong for assistance with atomic model building, and D. Silva-Sanchez for assistance with CryoJax analysis.

## Author Contributions

A.C.C. and G.M.A. conceived of the study. A.C.C. performed all cryo-EM and biochemical experiments. M.J.R. developed the neural network-based particle picker. F.M. and G.M.H. performed and analyzed MD simulations. A.C.C. and G.M.A. wrote the paper with input from all authors.

## Data Availability

Cryo-EM density maps and atomic models have been deposited in the EMDB and PDB with the following accession codes: K255E ACTN4 bound to F-actin [EMDB EMD-XXXX, PDB XXXX]; Wild-type ACTN4 bound to F-actin (strong state) [EMDB EMD-XXXX]; Wild-type ACTN4 bound to F-actin (weak state) [EMDB EMD-XXXX]; Wild-type ACTN4 bound to F-actin under force (+ATP data only) [EMDB EMD-XXXX]. Trained neural networks used for cryo-EM micrograph denoising and segmentation are available on Zenodo: https://doi.org/10.5281/zenodo.18841642. Simulation results are available at: https://github.com/hocky-research-group/Chin_Mukadum_Actinin4_simulation_2026. All other data are presented in the manuscript. Materials and reagents are freely available from the corresponding author (G.M.A.) upon request.

## Code Availability

Custom code for cryo-EM micrograph denoising and segmentation is available at: https://www.github.com/alushinlab/ACTN4. Simulation analysis code is available at: https://github.com/hocky-research-group/Chin_Mukadum_Actinin4_simulation_2026.

## Materials and Methods

### Plasmids and cloning

Polymerase Chain Reactions (PCR) were performed using KOD Hot Start Master Mix (MilliporeSigma, 71842) or Q5 Hot Start High-Fidelity 2X Master Mix (New England Biolabs, M0494L), and PCR products were purified using a QIAquick Gel Extraction Kit (Qiagen, 28704). Plasmid assembly and site-directed mutagenesis were performed using Gibson assembly (New England Biolabs, E2611L) or kinase-ligase-DpnI (KLD) cloning (New England Biolabs, M0554S). All plasmids were cloned and propagated in NEB 5-alpha competent *E. coli* cells (New England Biolabs, C2987U) and purified using a QIAprep Spin Miniprep Kit (Qiagen, 27106) or PureYield Plasmid Maxiprep System (Promega, A2393). All plasmids were verified by whole plasmid sequencing (Plasmidsaurus).

pET bacterial expression vectors encoding N-terminally 6xHis-SUMO tagged human ACTN4 and its variants (Fig. S1A) were prepared. For fluorescent labelling, a construct was prepared with HaloTag C-terminal to ACTN4. Previously reported pET bacterial expression vectors encoding N-terminally 6xHis tagged Ulp1^41^ and anti-GFP nanobody LaG94-10^64^ were employed in this study, as were pCAG mammalian expression vectors encoding C-terminally GFP-Flag tagged human myosin Va (AA 1−1091) and untagged human calmodulin (AA 1−149)^65^. A plasmid for mammalian cell expression of mature human α-actin-1 (AA 3−377) with an N-terminal 6xHis-Flag tag and traceless TEV protease cleavage site was purchased^66^ (Addgene, 188452); site-directed mutagenesis was performed on this plasmid to create ΔDEDE, D26K/D27K, and E101K/E102K variants.

### SDS-PAGE

Samples were run on NuPAGE 12% Bis-Tris gels (Invitrogen, NP0343BOX) using NuPAGE MOPS SDS Running Buffer (Invitrogen, NP000102), stained with Coomassie Brilliant Blue R-250 (Thermo Scientific, 20278), and imaged (Bio-Rad GelDoc Go). Band intensities were quantified using Fiji^67^.

### Recombinant bacterial protein expression and purification

All ACTN4 proteins were expressed in Rosetta 2(DE3) Singles Competent Cells (MilliporeSigma, 71400) in a 10 ml LB (MilliporeSigma, L3022) starter culture containing ampicillin (100 µg/mL) overnight at 37°C and added to 1 L LB containing ampicillin and chloramphenicol (25 µg/mL), which was grown at 37°C to OD600 0.6, induced with 0.5 mM IPTG, and grown for 6 hours at 37°C. The cells were pelleted, resuspended with ice-cold DPBS (Gibco, 14190144), pelleted, snap frozen in liquid nitrogen, and stored at −80°C. All subsequent purification steps were performed at 4°C. Each cell pellet was resuspended in 35 ml lysis buffer containing 50 mM Tris-HCl pH 8, 150 mM NaCl, 20 mM imidazole, 5% v/v glycerol, 2 mM dithiothreitol (DTT), 1 mM phenylmethylsulfonyl fluoride (PMSF), and 1:200 Protease Inhibitor Cocktail Set III EDTA-Free (Calbiochem, 539134) and high-pressure homogenized (Avestin Emulsiflex C5). Lysate was clarified at 31,000 × g for 30 minutes and incubated with 1.5 ml HisPur Cobalt Resin (Thermo Fisher Scientific, 89964) pre-equilibrated with lysis buffer for 2.5 hours on a rotator. After loading the sample onto a gravity flow column (Bio-Rad, 7321010) and discarding flow-through, the resin was washed with 66 bed volumes of wash buffer (lysis buffer without DTT, PMSF, or protease inhibitor). The resin was resuspended in 4 bed volumes of elution buffer (wash buffer but at 100 mM imidazole) for 30 minutes. After adding 0.85 mg 6xHis-Ulp1, the eluate was dialyzed against a buffer containing 50 mM Tris-HCl pH 8.0, 20 mM NaCl, and 1 mM DTT overnight in a 10K MWCO SnakeSkin dialysis membrane (Thermo Fisher Scientific, 68100).

The protein was then reapplied to the same resin, washed with dialysis buffer, to bind 6xHis-Ulp1 and uncleaved ACTN4, and the eluate was concentrated by a 50K Nominal Molecular Weight Limit (NMWL) centrifugal filter (MilliporeSigma, UFC805024) and clarified at 21,000 × g by tabletop centrifuge for 10 minutes. The protein was loaded onto a HiTrap Q HP 5 mL (Cytiva, 17115401) anion exchange column equilibrated with 50 mM Tris-HCl pH 8, 20 mM NaCl, and 1 mM DTT and eluted with a linear gradient to 1 M NaCl. Pooled fractions verified by SDS-PAGE were concentrated by a 50K NMWL centrifugal filter and loaded onto a Superdex 200 Increase 10/300 GL (Cytiva, 28-9909-44) size exclusion column equilibrated with 20 mM Tris-HCl pH 8, 100 mM NaCl, and 1 mM DTT. Pooled fractions verified by SDS-PAGE were concentrated by a 50K NMWL centrifugal filter, aliquoted, snap frozen in liquid nitrogen, and stored at −80°C. Notably, unlike all ACTN4 constructs that included the N-terminal disordered region, WT^ΔNterm^ ACTN4 was inefficiently cleaved by 6xHis-Ulp1. Uncleaved WT^ΔNterm^ ACTN4 was cleared, as verified by disappearance of a heavier band on SDS-PAGE, after size exclusion chromatography by incubation with 1 ml of Ni-NTA Agarose (Qiagen, 30210) pre-equilibrated with size exclusion buffer for 1 hour on a rotator.

6xHis-Ulp1 was expressed in BL21(DE3) Competent *E. coli* cells (New England Biolabs, C2527H) in a 10 ml LB starter culture containing kanamycin (50 µg/mL) overnight at 37°C and added to 1 L LB containing kanamycin, which was grown at 37°C to OD600 0.8, induced with 0.7 mM IPTG, and grown for 16 hours at 16°C. The cells were pelleted, resuspended with ice-cold DPBS, pelleted, snap frozen in liquid nitrogen, and stored at −80°C. All subsequent purification steps were performed at 4°C. Each cell pellet was resuspended in 35 ml lysis buffer (same as that for ACTN4 but without PMSF or protease inhibitor) and high-pressure homogenized. Lysate was clarified at 31,000 × g for 30 minutes and incubated with 1.5 ml Ni-NTA Agarose pre-equilibrated with lysis buffer for 1 hour on a rotator. After loading the sample onto a gravity flow column and discarding flow-through, the resin was washed with 66 bed volumes of wash buffer (same as that for ACTN4). The resin was resuspended in 4 bed volumes of elution buffer (wash buffer but including 300 mM imidazole) for 30 minutes. The eluate was dialyzed against a storage buffer containing 50 mM Tris-HCl pH 8, 200 mM NaCl, 1 mM DTT, and 20% v/v glycerol overnight in a 10K MWCO SnakeSkin dialysis membrane. After verification by SDS-PAGE, the protein was aliquoted, snap frozen in liquid nitrogen, and stored at −80°C.

### Skeletal muscle actin purification

G-actin was prepared based on a previously described protocol^68^. All purification steps were performed at 4°C. Chicken skeletal muscle acetone powder was resuspended in G-Ca buffer containing 2 mM Tris-HCl pH 8, 0.5 mM DTT, 0.2 mM ATP, 0.1 mM CaCl_2_, and 0.01% NaN_3_ (18 ml buffer per 1 g powder) for 30 minutes on a rotator. After ultracentrifugation in a Beckman Coulter Ti70 rotor at 42,500 rpm for 30 minutes, the supernatant was filtered through filter paper (Whatman, 1002-090). While stirring, 50 mM KCl and 2 mM MgCl_2_ were added to initiate actin polymerization. After 1 hour, 0.8 M KCl was added to dissociate proteins bound to F-actin. After 30 minutes, the solution was ultracentrifuged in a Beckman Coulter Ti70 rotor at 42,500 rpm for 3 hours. The supernatant was discarded, and approximately 2 ml of G-Ca buffer was added to the pellet, ensuring it was fully submerged, and incubated overnight. The mixture was Dounce homogenized, sheared through 26G then 30G needles, and dialyzed into G-Ca buffer overnight via a 10K MWCO SnakeSkin dialysis membrane. The sample was sheared again through a 30G needle and dialyzed again into G-Ca buffer overnight via a 10K MWCO SnakeSkin dialysis membrane. After ultracentrifugation in a Beckman Coulter Ti90 rotor at 70,000 rpm for 3 hours, the upper 2/3 of supernatant was loaded onto a HiLoad 16/600 Superdex 200 pg (Cytiva, 28-9893-35) size exclusion column equilibrated with G-Ca buffer containing 0.5 mM tris(2-carboxyethyl)phosphine hydrochloride (TCEP) instead of 0.5 mM DTT and set to detect absorbance at 290 nm. The second half of the peak was collected, verified by SDS-PAGE, and stored at 4°C. Unless otherwise noted, experiments were performed using actin purified from chicken skeletal muscle.

### Recombinant mammalian protein expression and purification

Myosin V-GFP and recombinant actins were expressed in FreeStyle 293-F cells (Thermo Fisher Scientific, R79007) cultured in FreeStyle 293 expression medium (Thermo Fisher Scientific, 12338026) at 37°C and 8% CO_2_. Per 400 ml of cell culture at 1.5 × 10^6^ cells/ml density, 400 μg of plasmid (6:1 mass ratio of myosin to calmodulin was co-transfected) was mixed with 1.2 ml of 1 mg/ml PEI MAX (Polysciences, 24765) in 15 ml medium. The mixture was shaken briefly and incubated for 20 minutes at room temperature prior to co-transfection. After 72 hours, the cells were pelleted, resuspended with ice-cold DPBS, pelleted, snap frozen in liquid nitrogen, and stored at −80°C. All subsequent purification steps were performed at 4°C.

Myosin V-GFP cell pellets from 720 ml of culture were resuspended in 30 ml Myosin Lysis Buffer containing 50 mM Tris-HCl pH 7.5, 150 mM NaCl, 2 mM MgCl_2_, 2 mM ATP, 0.2% 3-((3-cholamidopropyl) dimethylammonio)-1-propanesulfonate (CHAPS), 1 μg/ml aprotinin, 1 μg/ml pepstatin A, 1 μg/ml leupeptin, and 1 mM PMSF for 1 hour on a rotator. Lysate was clarified at 27,000 × g for 30 minutes and incubated with 400 μl of anti-Flag M2 affinity gel (MilliporeSigma, A2220), which was prewashed with 0.1 M glycine HCl pH 3.5 and equilibrated in Myosin Wash Buffer (Lysis Buffer without CHAPS, aprotinin, pepstatin A, leupeptin, or PMSF), for 1.5 hours on a rotator. After spinning down the gel and decanting the supernatant, the gel was resuspended in Myosin Lysis Buffer, transferred to a microtube, and washed with Myosin Wash Buffer three times. The gel was then incubated with 100 μg/ml Flag peptide (MilliporeSigma, F3290) in Myosin Wash Buffer on a rotator three times to elute myosin V-GFP. After buffer exchanging into a storage buffer containing 10 mM Tris-HCl pH 7.5, 100 mM NaCl, 2 mM MgCl_2_, and 3 mM DTT by concentrating with a 50K NMWL centrifugal filter, myosin V-GFP was aliquoted, snap frozen in liquid nitrogen, and stored at −80°C.

Recombinant actin cell pellets from 800 ml of culture were resuspended in 35 ml Actin Lysis Buffer containing 10 mM Tris-HCl pH 8, 10 mM CaCl_2_, 2 mM MgCl_2_, 0.5 mM ATP, 0.2% CHAPS, 10 μg/ml aprotinin, 10 μg/ml pepstatin A, 10 μg/ml leupeptin, and 2 mM PMSF for 3 hours on a rotator. Lysate was clarified at 50,000 × g for 20 minutes and incubated with 250 μl of DYKDDDDK Fab-Trap Agarose (Chromotek, ffa) pre-equilibrated with Actin Lysis Buffer for 2 hours on a rotator. After spinning down the Flag Fab-Trap resin and decanting the supernatant, the resin was washed with Actin Lysis Buffer three times, resuspended in Actin Lysis Buffer, transferred to a microtube, and washed with Actin Wash Buffer (lysis buffer without CHAPS, aprotinin, pepstatin A, leupeptin, or PMSF but supplemented with 1 mM DTT) three times. The resin was then incubated with a 1 ml solution comprised of 910 μl Actin Wash Buffer, 40 μl reconstituted Numacut TEV Protease 6x His-Tag (Numaferm, P1065-3), and 50 μl of 20x Numacut reaction buffer (supplied with the protease) overnight on a rotator to tracelessly release untagged recombinant actin into solution. After spinning down the resin, the supernatant was incubated with 270 μl Ni-NTA Agarose, equilibrated with wash buffer, in a new microtube for 2 hours on a rotator to pull down the protease and uncleaved recombinant actin. After spinning down the Ni-NTA Agarose, the supernatant was concentrated by a 30K NMWL centrifugal filter (MilliporeSigma, UFC803096), clarified at 21,000 × g by tabletop centrifuge for 10 minutes, and loaded onto a Superdex 200 Increase 10/300 GL size exclusion column equilibrated with Actin Wash Buffer. Pooled fractions verified by SDS-PAGE were concentrated by a 30K NMWL centrifugal filter and stored at 4°C.

### Preparation of anti-GFP nanobody coupled to fibrinogen

Anti-GFP nanobody-fibrinogen was prepared based on previously described protocols^64,69^. Anti-GFP nanobody was expressed in BL21(DE3) Competent *E. coli* cells in a 10 ml LB starter culture containing ampicillin (100 µg/mL) overnight at 37°C and added to 1 L LB containing ampicillin (100 µg/mL), which was grown at 37°C to OD600 0.4, induced with 0.1 mM IPTG, and grown for 18 hours at 12°C. The cells were pelleted, resuspended with ice-cold DPBS, pelleted, snap frozen in liquid nitrogen, and stored at −80°C. All subsequent purification and conjugation steps were performed at 4°C unless otherwise noted. Each cell pellet was resuspended in 10 ml TES buffer (0.2 M Tris-HCl pH 8, 0.5 mM EDTA, 0.5 M sucrose) and subjected to osmotic shock for 30 minutes by addition of 15 ml of TES buffer diluted 1:4 with water. The soluble periplasmic fraction was isolated by clarification at 4,000 × g for 10 minutes followed by another clarification at 48,000 × g for 30 minutes. The supernatant was mixed with an equal volume of Nanobody Wash Buffer containing 20 mM sodium phosphate pH 8.0, 1 M NaCl, and 20 mM imidazole and incubated with 2 ml Ni-NTA Agarose pre-equilibrated with Nanobdy Wash Buffer for 20 minutes on a rotator. After spinning down the resin and discarding the supernatant, the resin was incubated with 50 ml of Nanobody Wash Buffer for 20 minutes on a rotator. After loading the sample onto a gravity flow column and discarding flow-through, the protein was eluted in 1 ml fractions with Nanobody Elution Buffer containing 20 mM sodium phosphate pH 8, 0.5 M NaCl, and 0.3 M imidazole. Pooled fractions verified by SDS-PAGE were concentrated by a 3K NMWL centrifugal filter (MilliporeSigma, UFC800396) and buffer exchanged into Fibrinogen Buffer (0.1 M sodium bicarbonate and 0.5 mM EDTA pH 8.3) supplemented with 1 mM TCEP.

Fibrinogen (MilliporeSigma, 341576-M) was resuspended at 4 mg/ml in Fibrinogen Buffer and incubated with 25-fold molar excess of maleimide-PEG8-succinimidyl ester (MilliporeSigma, 746207) for 1 hour at room temperature on a rocker. Excess maleimide-PEG8-succinimidyl ester was cleared via a Zeba Spin Desalting Column 7K MWCO equilibrated in Fibrinogen Buffer. TCEP was removed from the anti-GFP nanobody via a Zeba Spin Desalting Column 7K MWCO pre-equilibrated in Fibrinogen Buffer. Fibrinogen-maleimide-PEG8-succinimidyl ester was incubated with five-fold molar excess of anti-GFP nanobody overnight on a rotator. To quench unreacted maleimide-PEG8-succinimidyl ester, 50 mM cysteine was added and incubated for 30 minutes at room temperature. The sample was brought to 25% final ammonium sulfate and incubated for 30 minutes at room temperature on a rocker to precipitate the anti-GFP nanobody-fibrinogen conjugate. After pelleting at 16,873 × g for 10 minutes at room temperature, followed by resuspension in the initial volume of Fibrinogen Buffer, a second round of precipitation with ammonium sulfate and pelleting was performed. The sample was resuspended to 5 mg/ml, aliquoted, snap frozen in liquid nitrogen, and stored at −80°C.

### Preparation of Lifeact peptide

Lifeact (MGVADLIKKFESISKEE) was generated by chemical peptide synthesis (GenScript) with HPLC purity of 99.2% and verified by LC-MS, NMR, and qualitative solubility test in DPBS.

### Cryo-EM grid preparation (force-free conditions)

C-flat Holey Carbon Grids (Gold 1.2 µm hole 1.3 µm space 300 mesh, Electron Microscopy Sciences, CF313-50-Au) were plasma cleaned with a hydrogen and oxygen mixture for 15 seconds (Gatan Solarus Model 950 Advanced Plasma System). F-actin was prepared by polymerizing G-actin with 2 molar equivalents of phalloidin (Thermo Fisher Scientific, P3457), KMEI (50 mM KCl, 1 mM MgCl_2_, 1 mM EGTA, 10 mM imidazole pH 7), and G-Mg (2 mM Tris-HCl pH 8, 0.5 mM DTT, 0.2 mM ATP, 0.01% NaN_3_, 0.1 mM MgCl_2_) supplemented with 0.01% Nonidet P40 Substitute (MilliporeSigma, 11332473001) for 1 hour at room temperature followed by incubation overnight at 4°C. K255E ACTN4 was incubated with F-actin for 1 hour at 4°C at a 1:8 ratio (0.5 µM or 0.2 µM F-actin) immediately before freezing grids. A pilot K255E ACTN4 dataset (Fig. S4D) was prepared similarly except at a 1:4 ratio (0.5 µM F-actin). Wild-type ACTN4 samples were prepared by overnight 4°C incubation at a 1:4 ratio (0.2 µM or 0.1 µM F-actin). In the humidified (90%) chamber of a Leica EM GP plunge freezer operating at 25°C, 4 µl of ACTN4/F-actin mixture was applied to the grid for 1 minute. The grid was then back-blotted for 4 seconds, plunge frozen in liquid ethane, and transferred to liquid nitrogen for storage.

### Force reconstitution on cryo-EM grids

F-actin was prepared by copolymerizing 0.5 μM G-actin, 2 molar equivalents of Alexa Fluor 568 phalloidin (Thermo Fisher Scientific, A12380), KMEI, and G-Mg for 1 hour at room temperature, then kept on ice. All subsequent samples were prepared on ice. Myosin V-GFP and anti-GFP nanobody-fibrinogen were diluted in Motility Buffer (20 mM 3-(N-morpholino)propanesulfonic acid (MOPS) pH 7.4, 5 mM MgCl_2_, 0.1 mM EGTA, 50 mM KCl, 1 mM DTT) to 0.5 µM and 0.5 mg/ml respectively. Myosin V-GFP preparations were supplemented with 10 U/ml apyrase (New England Biolabs, M0398S) for −ATP conditions. Blocking Buffer was prepared with 0.01 mg/ml PLL(20)-g[3.5]-PEG(2) (SuSoS), 0.2 mg/ml κ-casein (MilliporeSigma, C0406), and 1% polyvinylpyrrolidone (MilliporeSigma, PVP10). Wild-type ACTN4-Halo was labeled with 2 molar equivalents of Janelia Fluor 646 HaloTag Ligand (Promega, GA1120) in Motility Buffer for 2 hours on a rotator at 4°C, cleared of excess dye via a Zeba Spin Desalting Column 7K MWCO (Thermo Fisher Scientific, 89882), equilibrated with Motility Buffer, and diluted to 20 nM. Imaging Buffer was prepared with 0.1 mg/ml glucose oxidase (MilliporeSigma, G7141), 0.02 mg/ml catalase (MilliporeSigma, C3515), 2 molar equivalents (to actin) of unlabeled phalloidin, in Motility Buffer supplemented with 15 mM glucose. Samples were prepared in the presence and absence of 20 nM labelled wild-type ACTN4-Halo (which was omitted for experiments displayed in Fig. 3B) and 2 mM ATP (which was omitted for −ATP conditions). All reagents except F-actin were prepared in 0.2 ml open-top thickwall polycarbonate tubes 7 × 20 mm (Beckman Coulter, 343775) and clarified by ultracentrifugation in a Beckman Coulter TLA-100 rotor at 100,000 × g for 12 minutes at 4°C.

C-flat Holey Carbon Grids (Gold 4.0 µm hole 2.0 µm space 300 Mesh, Electron Microscopy Sciences, CF423-50-Au) were plasma cleaned with a hydrogen and oxygen mixture for 15 seconds. Wells were prepared by attaching a PDMS Stencil (Alvéole, circle wells of Ø 4mm) to a No. 1.5 24 × 60 mm glass coverslip (Corning, 2980-246). Each grid was treated in the wells in the following sequence: anti-GFP nanobody-fibrinogen (3 minutes), Blocking Buffer (1 minute), myosin V-GFP (2 minutes), F-actin (4 minutes), and labelled wild-type ACTN4-Halo (2 minutes; a buffer blank was used for experiments in Fig. 3B. Wells were washed with Motility Buffer after Blocking Buffer, myosin V-GFP, and F-actin steps. Each grid was removed from the well, submerged (carbon facing down) in a few drops of imaging buffer on the glass slide, and sandwiched by a No. 1.5 22 × 22 mm glass coverslip (Fisherbrand, 12-541-B).

Time-lapse dual-color epifluorescence imaging was performed using a Nikon Eclipse Ti-E microscope illuminated by 561 nm (50 mW) and 640 nm (45 mW) lasers from an Agilent MLC400B Monolithic Laser Combiner. Using Nikon NIS-Elements software, images were acquired either every 1 second or 2 seconds through a CFI Apochromat TIRF 100XC Oil objective (used in combination with a 1.5× optivar), quad filter (Chroma), and iXon 897 EMCCD camera (Andor). F-actin bundle curvature was measured using the Kappa plugin in Fiji^67^.

Grids for cryo-fluorescence and cryo-EM were prepared similarly except using 0.8 μM G-actin copolymerized with Alexa Fluor Plus 555 phalloidin (Thermo Fisher Scientific, A30106), untagged wild-type ACTN4, C-Flat Holey Carbon Grid Gold 2.0 µm hole 1.0 µm space 300 mesh (Electron Microscopy Sciences, CF312-50-Au), and Cryo-EM Motility Buffer (Imaging Buffer without glucose oxidase, catalase, or glucose but supplemented with 0.01% Nonidet P40 substitute) instead of Imaging Buffer. After treatment with Cryo-EM Motility Buffer for 2 minutes, grids were back-blotted for 4 seconds (with 5 seconds post-blot time), plunge frozen in liquid ethane, and transferred to liquid nitrogen for storage.

Based on our preparation procedure, the F-actin in our force reconstitution assays was in a phalloidin-stabilized, ADP-P_i_ state, which rigidifies the filament^40^. We used this highly stable form of F-actin to facilitate transmission of load to the ACTN4 crosslinker through F-actin while minimizing force-induced filament ruptures. For consistency, the F-actin used in our cryo-EM structures under force-free conditions was also in this phalloidin-stabilized, ADP-P_i_ state.

### Cryo-fluorescence imaging

Grids were mounted on a Leica THUNDER EM Cryo CLEM microscope cooled to −195°C maintained in a room with relative humidity below 30%. Epifluorescence microscopy was performed using a 50× 0.9 NA air objective. Brightfield and rhodamine (EX: 541-551, DC: 560, EM: 565-605) channels were used with 1 ms and 200 ms exposure times, respectively. Z-stacks of images were manually set around a focus point using system optimized settings and captured at a depth of 16-bit. Images were acquired using LAS X software and postprocessed with THUNDER Small Volume Computational Clearing.

### Cryo-EM data collection

Datasets were collected on a FEI Titan Krios system operating at 300 kV and equipped with a Gatan K3 detector and GIF BioQuantum energy filter (slit width of 20 eV) using super-resolution mode. Movies were recorded using the SerialEM software suite^70^ at a nominal magnification of ×81,000, corresponding to a calibrated pixel size of 1.09 Å at the specimen level (super-resolution pixel size of 0.545 Å/pixel). Each 4.5 seconds exposure was dose-fractionated across 50 frames, with a total electron dose of 49.238 e^−^/Å^2^ (0.9848 e^−^/Å^2^/frame) over a defocus range of −0.8 to −2.2 µm. K255E ACTN4–F-actin datasets were collected at 30°, 15°, and 0° tilts using 1 shot per hole through stage translations. Wild-type ACTN4–F-actin datasets were collected at 0° tilt, using 1 shot per hole for data collected in the absence of motors, and 3 shots per hole for data collected in the presence of motors. Beam-image shift was used to collect 0° tilt data from 9 holes (a 3×3 matrix) per stage translation.

The pilot K255E ACTN4 dataset used to generate a preliminary reconstruction was collected on a FEI Titan Krios system operating at 300 kV and equipped with a Gatan K2 Summit detector using super-resolution mode. Movies were recorded using the SerialEM software suite at a nominal magnification of ×29,000, corresponding to a calibrated pixel size of 1.03 Å at the specimen level (super-resolution pixel size of 0.515 Å/pixel). Each 10 seconds exposure was dose-fractionated across 40 frames with a total electron dose of 61.268 e−/Å^2^ (1.5317 e−/Å^2^/frame) over a defocus range of −0.8 to −2.2 µm. Data were collected at 30°, 15°, and 0° tilts using three shots per hole, and beam-image shift was used to collect 0° tilt data from 9 holes (a 3×3 matrix) per stage translation.

### Cryo-EM image pre-processing

Movies were aligned using MotionCor2^71^ with 5×5 patches, and dose-weighted sums were generated from twofold binned frames with Fourier cropping to a pixel size of 1.09 Å. Contrast transfer function (CTF) parameter estimation was performed with CTFFIND4^72^ using non-dose-weighted sums. Due to significant heterogeneity in the density of F-actin filaments across grids in the presence of ACTN4 crosslinker, micrographs from force-free conditions were manually curated to select those with clearly visible ACTN4–F-actin interfaces. A total of 25,402 micrographs were collected for K255E ACTN4–F-actin, and a subset of 4,033 curated micrographs were initially processed to obtain a high-resolution map that was then used as an initial model for obtaining more particles from the rest of the micrographs using an identical workflow. A total of 46,468 micrographs were collected for wild-type ACTN4–F-actin in the absence of motors, and 15,954 curated micrographs were processed. Unlike micrographs from motor-free conditions, which were often overcrowded, most micrographs from force reconstitution assays of wild-type ACTN4–F-actin in the presence of motors (both −ATP and +ATP conditions) were empty. For this reason, and to avoid biases regarding what constitutes clear ACTN4–F-actin interfaces, all micrographs with any F-actin present were kept for picking. A total of 30,108 and 50,822 micrographs were collected for −ATP and +ATP conditions respectively, and 8,104 (+ATP dataset only) and 15,735 (combined) curated micrographs were processed.

### Neural network particle picking

The machine learning-based particle picking approach used in this study was adapted from our previous cryo-EM analysis of cytoskeletal crosslinkers^40,41^. To create *in silico* models for generating a synthetic dataset for neural network training (Fig. S2A), a cryo-EM map of E254K filamin ABD bound to F-actin (EMD-8918), segmented to only include one occupied site, was manually combined with a map of half of the rod domain proximal to the ABD in ACTN2 (PDB: 4D1E, residues 257-513) created using the pdb2mrc function from EMAN2^73^. A library of 15 models, each within a 256-pixel (4.36 Å/pixel) box, was created with the rod domain in different poses to capture a range of flexibility. Volumes were rotated about the psi and rot angles by a random, uniformly sampled value between 0° and 359°, and the tilt was randomly sampled from a Gaussian probability distribution centered at 90° with a standard deviation of 10°. Volumes were then randomly translated around the box by ±218 Å and projected along the z-axis to generate noiseless projections. Projection images included 0, 1, 2, or 3 filaments with probabilities of 0.15, 0.6, 0.2, and 0.05, respectively. Each filament had a probability of 0.5 to be bare F-actin and 0.033 (1/30) to be each of the 15 models. Paired noisy projections were generated by corrupting with a CTF then adding empirical pink noise in Fourier space. Projection images were cropped to a smaller box size of 128 pixels to ensure that filaments fully span the box. Three-channel stacks of semantic maps associated with noisy/noiseless projection pairs were generated by low-pass filtering the noiseless projection by 30 Å, binarizing it, and assigning it as background, bare F-actin, or ACTN4 bound F-actin.

Neural network architectures, loss functions, learning rates, minibatch sizes, and training/validation splits used in this study are identical to those described our previous study^41^. The denoising autoencoder (DAE) was trained using 200,000 noisy/noiseless projection pairs with box sizes of 128×128. Upon network convergence, the DAE had an average cross-correlation coefficient of 0.9908 on the validation set. The semantic segmentation network was trained using 10,000 pairs of noisy inputs and semantically segmented targets of dimensions 128×128 and 128×128×3, respectively. Upon network convergence, the semantic segmentation network had an average categorical cross-entropy of 0.1058 on the validation set.

To pick particles, motion-corrected micrographs were down-sampled by 4 and converted into semantic maps by extracting 128-pixel boxes across the micrograph in a raster pattern with 32 pixels of overlap and stitching the output into a semantic map by computing a maximum intensity projection of the overlapping regions. Only the decorated F-actin channel results were used (Fig. S2B), and they were binarized using a fixed threshold of 0.8. Binarized images were skeletonized, and non-maximum suppression was used to ensure all particle picks were spaced at least 44 Å apart. The particle coordinates were used for extraction.

### Cryo-EM data processing: motor-free specimens

The data processing workflow for K255E ACTN4–F-actin is displayed in Fig. S3A. Particles picked by the neural network were imported into RELION-4.0^74^ and extracted with a box size of 512 pixels, then binned by 8. Extracted particles were imported into cryoSPARC v4.2.1^75^ for reference-free 2D classification. Selected particles were used for *ab initio* reconstruction. Particles from filamentous classes were subjected to homogeneous refinement using an initial model from a pilot dataset, which was not curated. The preliminary map determined using this pilot dataset featured low-resolution CH1 and CH2 domain densities all along the filament (Fig. S4D). It was therefore segmented in UCSF Chimera^76^ to include only one CH1 domain density on F-actin and binned by 8 to produce an initial reference.

Particles were re-extracted at bin 4 in RELION-4.0 and subjected to 3D auto-refinement and 3D classification. Particles from the class with CH1 domain density were subjected to 3D auto-refinement, re-extraction at bin 2, masked 3D auto-refinement (with a mask covering F-actin and the CH1 domain), and masked 3D classification. After removing duplicate particles within 20 Å, particles from the class with CH1 domain density were subjected to masked 3D auto-refinement. The resultant map was used as an initial model for processing additional particles, picked by the neural network on micrographs that were not originally curated, using an identical workflow. Combining particles from both datasets, masked 3D auto-refinement and focused 3D classification (with a mask around CH1 domain and adjacent actin) were performed. Particles from the class with clear CH1 domain density were subjected to masked 3D auto-refinement, re-extraction at bin 1, masked 3D auto-refinement, and focused 3D classification. After masked 3D auto-refinement of particles from the class with clear CH1 domain density, iterative rounds of CTF refinement, Bayesian polishing, and masked 3D auto-refinement were performed. The resultant map was used as an initial model for processing the discarded micrographs from both datasets using an identical workflow. The combined particles were subjected to masked 3D auto-refinement and focused 3D classification. After removing duplicate particles within 27 Å (the approximate axial rise of F-actin), particles from classes with clear CH1 domain density were subjected to iterative rounds of CTF refinement, Bayesian polishing, and masked 3D auto-refinement. Final postprocessing produced a density map with 3.2 Å global resolution by the Fourier shell correlation (FSC) 0.143 criterion.

The data processing workflow for wild-type ACTN4–F-actin in the absence of motors is shown in Fig. S5B. Particles picked by the neural network were imported into RELION-4.0 and extracted with a box size of 320 pixels then binned by 8. Extracted particles were imported into cryoSPARC v4.2.1 for reference-free 2D classification. Particles from filamentous classes were subjected to masked homogeneous refinement (with a mask covering F-actin and the ABD) using the K255E ACTN4–F-actin map binned by 8 as an initial model, and unsupervised masked 3D classification was performed to discard particles that contributed to a distorted F-actin reconstruction. Masked homogeneous refinement and unsupervised focused 3D classification (with a mask covering the ABD and its adjacent actin) were performed, revealing putative weak and strong states. Particles from the weak state class were subjected to two rounds of masked homogeneous refinement and unsupervised focused 3D classification, and classes with clear ABD density were selected. The resultant map was used for supervised focused 3D classification to more effectively extract weak state particles from a second dataset, which was processed with a similar workflow to produce weak and strong state maps. Particles from the weak state maps of the first and second datasets were combined via masked homogeneous refinement, and this updated weak state map was used for supervised focused 3D classification to more effectively extract weak state particles from a third dataset, which was processed with a similar workflow to produce weak and strong state maps. Selected particle from all three datasets were then combined for further processing of each state.

To process the weak state, particles were re-extracted at bin 4 in RELION-4.0 and subjected to masked 3D auto-refinement and masked 3D classification. Particles from classes with clear ABD density were subjected to masked 3D auto-refinement and re-extracted at bin 2. After importing the particles into cryoSPARC v4.2.1, masked homogeneous refinement and unsupervised focused 3D classification were performed. Particles from the class with clear ABD density were subjected to masked homogeneous refinement and re-extracted at bin 1 in RELION- 4.0. After removing duplicate particles within 27 Å, masked 3D auto-refinement and CTF refinement were performed. Particles were imported into cryoSPARC v4.2.1 for masked 3D variability analysis. Particles from clusters with clear ABD density were subjected to masked local refinement and imported into RELION-4.0 for re-extraction at bin 1 with a box size of 512 pixels, Bayesian polishing, and masked 3D auto-refinement.

To process the strong state, particles were subjected to two rounds of supervised focused 3D classification, using a K255E ACTN4/F-actin map binned by 8 as an initial model, and masked homogeneous refinement. Particles from classes with clear ABD density were re-extracted at bin 4 in RELION-4.0 and imported into cryoSPARC v4.2.1 for masked local refinement and unsupervised focused 3D classification. Particles from classes with clear ABD density were subjected to masked local refinement and supervised focused 3D classification using a K255E ACTN4–F-actin map binned by 4 as an initial model. Particles from classes with clear ABD density were subjected to masked local refinement and re-extracted at bin 2 in RELION-4.0. After importing the particles into cryoSPARC v4.2.1, masked local refinement and supervised focused 3D classification, using a K255E ACTN4–F-actin map binned by 2 as an initial model, were performed. Particles from the class with clear ABD density were subjected to masked local refinement and unsupervised focused 3D classification. After masked local refinement of particles from the class with clear ABD density and re-extraction at bin 1 with a box size of 512 pixels in RELION-4.0, removal of duplicate particles within 27 Å, masked 3D auto-refinement, CTF refinement, and Bayesian polishing were performed. Final postprocessing produced density maps at a global resolution of 3.9 Å and 4.1 Å, for the weak state and the strong state, respectively, by the FSC 0.143 criterion, primarily driven by the well-resolved region of the density maps corresponding to F-actin. All 3D classification jobs performed in RELION-4.0 were without image alignment, and local resolution estimation was performed using RELION-4.0.

### Cryo-EM data processing: force-reconstitution specimens

Particles were picked by the neural network, extraction steps were performed with RELION-4.0, and all other processing steps were performed with cryoSPARC v4.2.1 / v.4.7.0. The data processing workflow for wild-type ACTN4-F-actin in the presence of motor forces (+ATP dataset only) is displayed in Fig. S10A. Particles were initially extracted with a box size of 512 pixels, binned by 4, and subjected to reference-free 2D classification. Selected particles were used for *ab initio* reconstruction, and particles contributing to filamentous classes were subjected to masked homogeneous refinement using a K255E ACTN4–F-actin map binned by 4 as an initial reference, followed by supervised focused 3D classification using bound and unbound F-actin references to remove bare filament segments. Particles from the class with ABD density were subjected to another round of masked homogeneous refinement and supervised focused 3D classification. Particles from classes with ABD density resembling the strong state were subjected to masked homogeneous refinement, then re-extracted at bin 2. Three rounds of masked local refinement and supervised focused 3D classification were performed. Particles from the class with clear ABD density were subjected to masked local refinement and re-extracted at bin 1. After removing duplicate particles within 27 Å, masked local refinement and supervised focused 3D classification were performed. Particles from the class with clear ABD density were subjected to a final masked local refinement, producing a of 6.7 Å resolution density map by the FSC 0.143 criterion.

The data processing workflow for wild-type ACTN4–F-actin under force (−ATP and +ATP datasets combined) is shown in Fig. S10B. Particles were extracted with a box size of 512 pixels, binned by 4, then subjected to reference-free 2D classification. Selected particles were used for *ab initio* reconstruction, followed by multiple rounds of masked local refinement and supervised focused 3D classification to remove segments that contained bound myosin V. To validate the identity of the bound ABP density, these particles were re-extracted at bin 1, duplicates particles within 27 Å were removed, followed by masked 3D auto-refinement. Final postprocessing produced a density map of 4.0 Å by the FSC 0.143 criterion with clear features of the myosin V motor domain.

The remaining particles were re-extracted with a box size of 320 pixels at bin 2, duplicate particles within 27 Å were removed, and a consensus masked local refinement was performed using a K255E ACTN4–F-actin map binned by 2 as an initial model, producing a cleaned particle stack for statistical analysis. Multiple rounds of supervised focused 3D classification and masked local refinement were performed using this particle stack to obtain maps corresponding to the strongly bound, weakly bound, and unbound states, which were used as references for supervised focused 3D classification. Using 6 classes (1 strong reference, 1 weak reference, and 4 unbound references) and a filter resolution of 15 Å, 10 independent runs of supervised focused 3D classification were performed. Particle Sets Tool in cryoSPARC was used to calculate the number of shared particles between volumes. For analysis of −ATP versus +ATP particles, csparc2star.py^77^ was used to extract metadata. Local resolution estimation was performed using RELION-4.0.

### Atomic model building

Previously determined atomic models of ADP-Pi-F-actin^40^ (PDB 8D14) and K255E ACTN4^34^ (PDB 2R0O; chain A, residues 47-154) were rigid-body docked into the cryo-EM density map in UCSF Chimera^76^. Residues 31-46 of K255E ACTN4 were built using Coot^78^. Structures were flexibly fitted with ISOLDE^79^. An atomic model of phalloidin from a previously determined structure^80^ (PDB 7PLU) was added to the flexibly fitted structures by rigid-body docking into cryo-EM density in UCSF Chimera. The final atomic model was produced through iterative rounds of real-space refine in Phenix^81^ and rebuilding in Coot, followed by a final round of real-space refinement. Model validation was conducted with MolProbity^82^ in Phenix.

### Subtilisin cleavage of actin

Subtilisin (type VIII protease from *B. licheniformis*; MilliporeSigma, P5380), reconstituted in DPBS, was added to G-actin at a mass ratio of 1:1000. After incubating at room temperature for 3 hours, the digestion was terminated with 2 mM PMSF.

### F-actin co-sedimentation assays

All samples were prepared at room temperature in microtubes. For assays using subtilisin-cleaved actin (Fig. S7D), F-actin was prepared by copolymerizing 4 μM G-actin, 5 molar equivalents of phalloidin, KMEI, and G-Mg for 3 hours before adding 0.4 μM ACTN4. For assays using recombinant actin (Fig. 2D; Fig. S7E), F-actin was prepared by copolymerizing 1 μM G-actin, 2 molar equivalents of phalloidin, KMEI, and G-Mg for 2 hours before adding 0.1 μM ACTN4. For all other assays, F-actin was prepared by copolymerizing 2 μM G-actin and 2 molar equivalents of phalloidin in KMEI and G-Mg for 1 hour before adding 0.2 µM ACTN4. For all assays, ACTN4 was incubated with F-actin for 30 minutes.

Samples were transferred to 0.2 ml 7 × 20 mm open-top thick wall polycarbonate tubes and ultracentrifuged in a Beckman Coulter TLA-100 rotor at 100,000 × g for 30 minutes at 4°C. Supernatant and pellet fractions were analyzed by SDS-PAGE.

To confirm that pelleting was due to F-actin binding, ACTN4 proteins thawed from snap frozen aliquots were ultracentrifuged under identical conditions but without actin (Fig. S1B). None of the ACTN4 proteins pelleted substantially with the exception of WT^ΔNterm^ and K255E^ΔNterm^ ACTN4. WT^ΔNterm^ ACTN4 was clarified at 100,000 × g for 30 minutes at 4°C prior to performing co-sedimentation assays, and the percent of residual pelleting was used to correct results obtained from co-sedimentation assays. K255E^ΔNterm^ ACTN4 completely pelleted and was therefore not used for co-sedimentation assays.

### Molecular dynamics simulations

Our starting atomic model used in simulations followed previously established protocols for simulating an actin filament^83^. We started with an equilibrated model of ADP-bound actin^52^ initiated from PDB model 2ZWH^84^ and selected three actin subunits to align with the proposed weak state models determined in our current study. Around these subunits, we created a solvation box with TIP3P water and 50 mM NaCl to neutralize the complex, mimicking physiological conditions. The resulting solvation box enclosing each system had dimensions of 11.5 x 13.5 x 15 nm^3^.

Minimization, equilibration, and production runs were performed with GROMACS-2023^85^. The CHARMM36^86^ force field was utilized for all the bonded and nonbonded interactions of the proteins. Each system was minimized with the steepest descent algorithm for 5000 steps with dt = 1, followed by a 5 ns constant-volume and temperature (NVT) equilibration stage, with a 2 fs timestep. We imposed position restraints on backbone heavy atoms (force constants = 1000 kJ/(mol * nm)) to allow the side chains to equilibrate with the solvent. Protein and solvent atoms were separately coupled to a 300K temp bath using the GROMACS v-rescale thermostat. Subsequently, we performed a 5 ns equilibration with a 2 fs timestep at constant temperature and pressure (NPT) using GROMACS’s V-Rescale thermostat and the C-Rescale barostat. For NPT simulations, only restraints on actin subunit a_i+1_ were retained, to preserve the orientation of the structure, while minimizing perturbations to the interactions between the ACTN4 ABD and its interacting actin subunits (a_i_ and a_i+2_). The resulting structure was then used for a 1 µs production run. This procedure was performed for 4 total orientations: CH1 PE, CH1 PE reverse face, CH2 PE, and CH2 PE reverse face. From the CH1 PE and CH2 PE starting runs, we performed 4 x 500 ns replica MD simulations initiated from time points from each microsecond simulation (50 ns, 100 ns, 150 ns, and 200 ns). The replica from each orientation that consistently showed the shortest CHD center-of-mass distance to actin was selected to run for an additional 1 µs. These runs were used to perform all subsequent map correlation analysis. For all equilibration and production runs, long-range electrostatics were calculated using the Particle Mesh Ewald (PME) algorithm with a 1.2 nm cut-off for short-range non-bonded interactions. Bonds between hydrogen and heavy atoms were constrained using the LINCS algorithm. Finally, the equations of motion were integrated every 2 fs using the Velocity-Verlet algorithm.

We used CryoJax^87^ to compare our simulations to the experimental cryo-EM map. The ridged-body docked model was used as a reference to align our simulations. We then generated simulated densities for each frame from a GROMACS trajectory file. Only non-hydrogen atoms from the ACTN4 ABD were included to avoid contributions from F-actin to overlap measurements. We additionally segmented ABD density from the cryo-EM map for comparison with simulated densities. A gaussian mixture atomic potential was generated on a (128,128,128) voxel grid, which was used for comparison with a down-sampled experimental map. The normalized cross-correlation between the simulated and real volumes was calculated by performing the dot product, normalized by the total root mean squared magnitudes of each volume, to return a value between 0 and 1, with 0 being no correlation between the densities and 1 having a perfect correlation.

### Molecular graphics and plotting

Cryo-EM maps and atomic models were visualized using UCSF Chimera^76^ and ChimeraX^88^. Plotting and statistical analysis were performed in Graphpad Prism.

---

**Figure S1:**
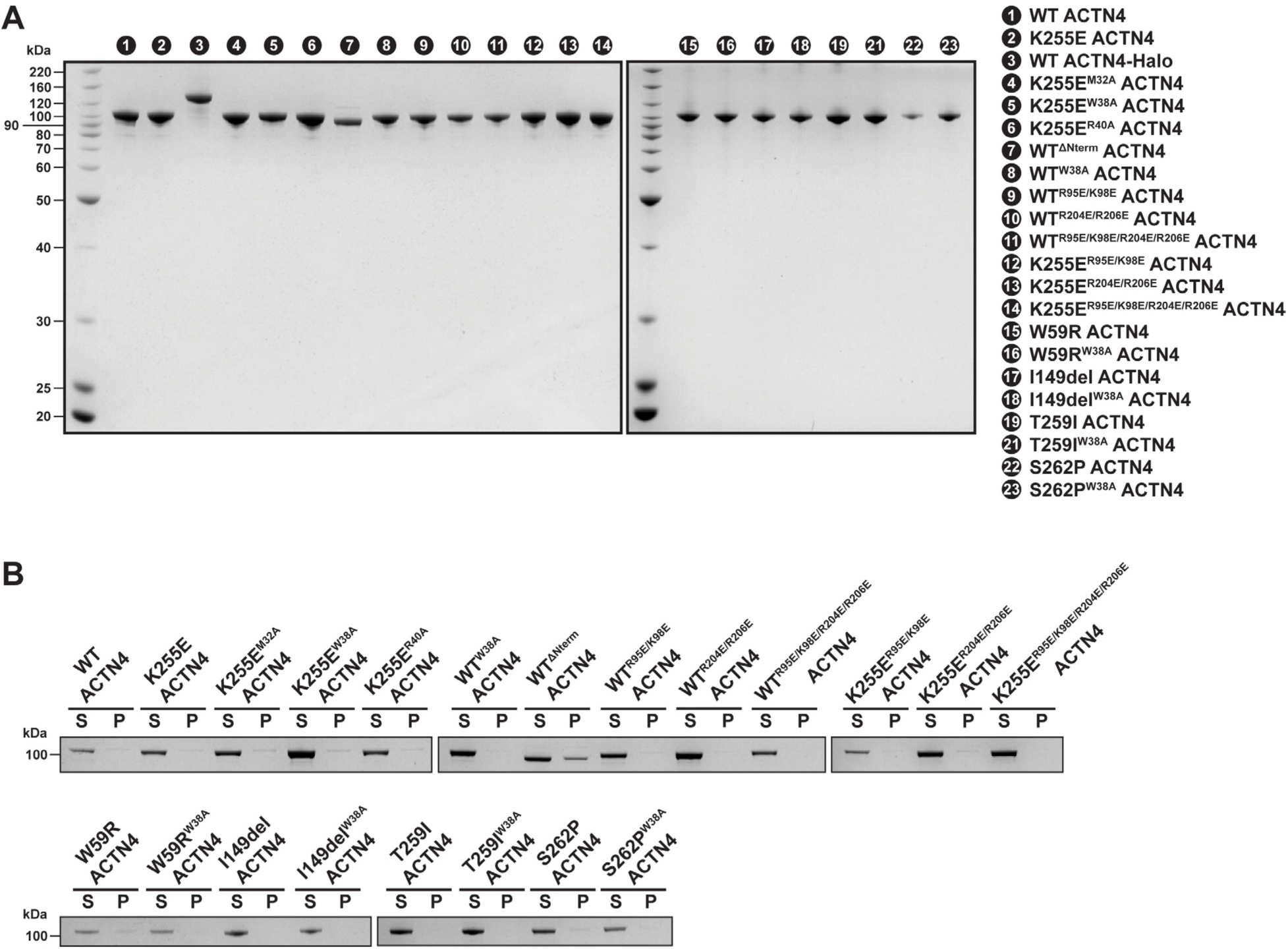
ACTN4 purification and sedimentation without actin. **(A)** SDS-PAGE of all purified ACTN4 proteins used in this study. **(B)** SDS-PAGE of control sedimentation assays performed in the absence of F-actin. WT^ΔN-term^ ACTN4 was the only construct which displayed substantial pelleting.

**Figure S2:**
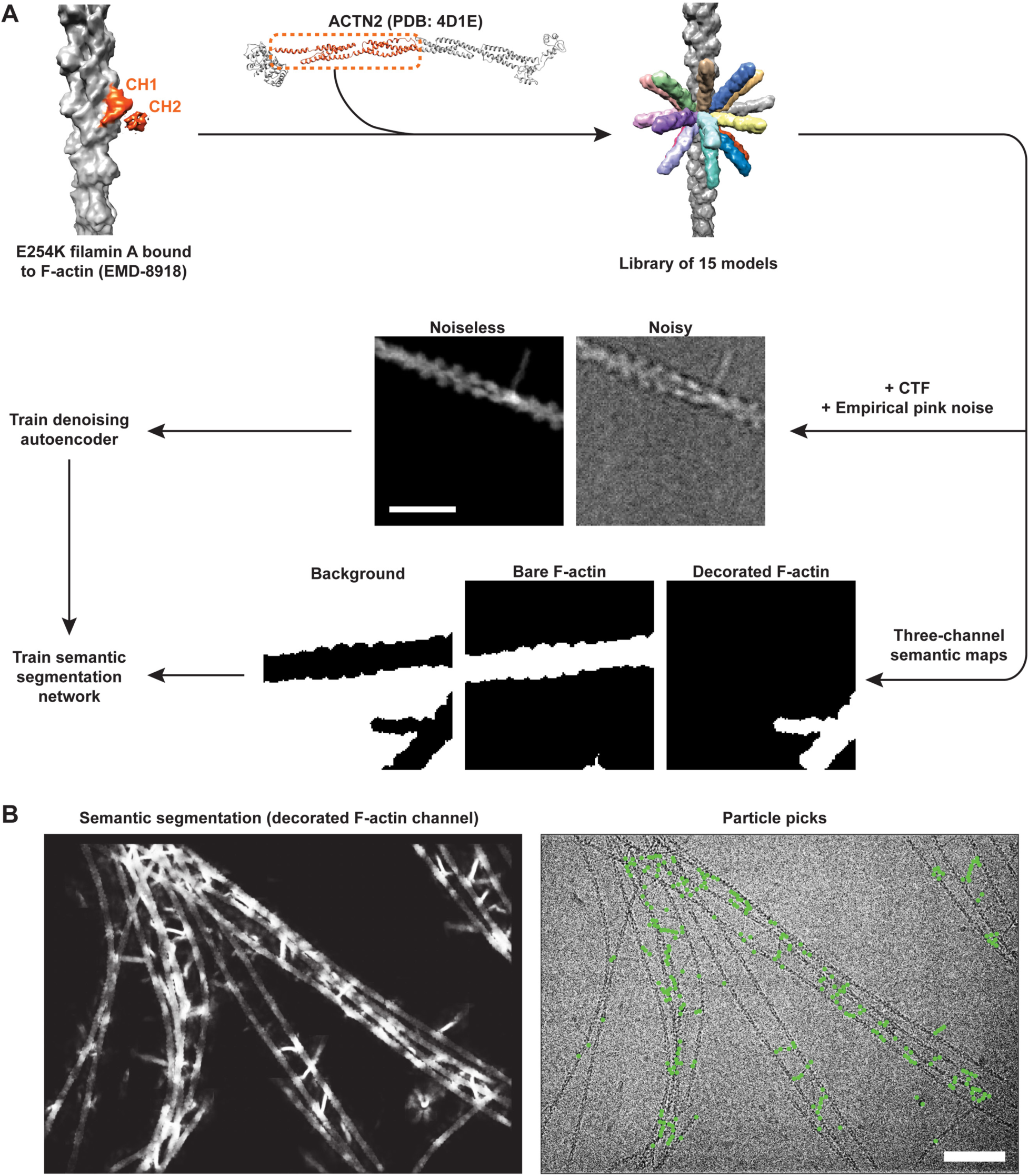
Neural network-based particle picker and example performance on experimental cryo-EM data. **(A)** Schematic of synthetic data generation and neural network training. Scale bar, 20 nm. **(B)** Network performance on a representative micrograph from the K255E ACTN4–F-actin dataset. Scale bar, 80 nm.

**Figure S3:**
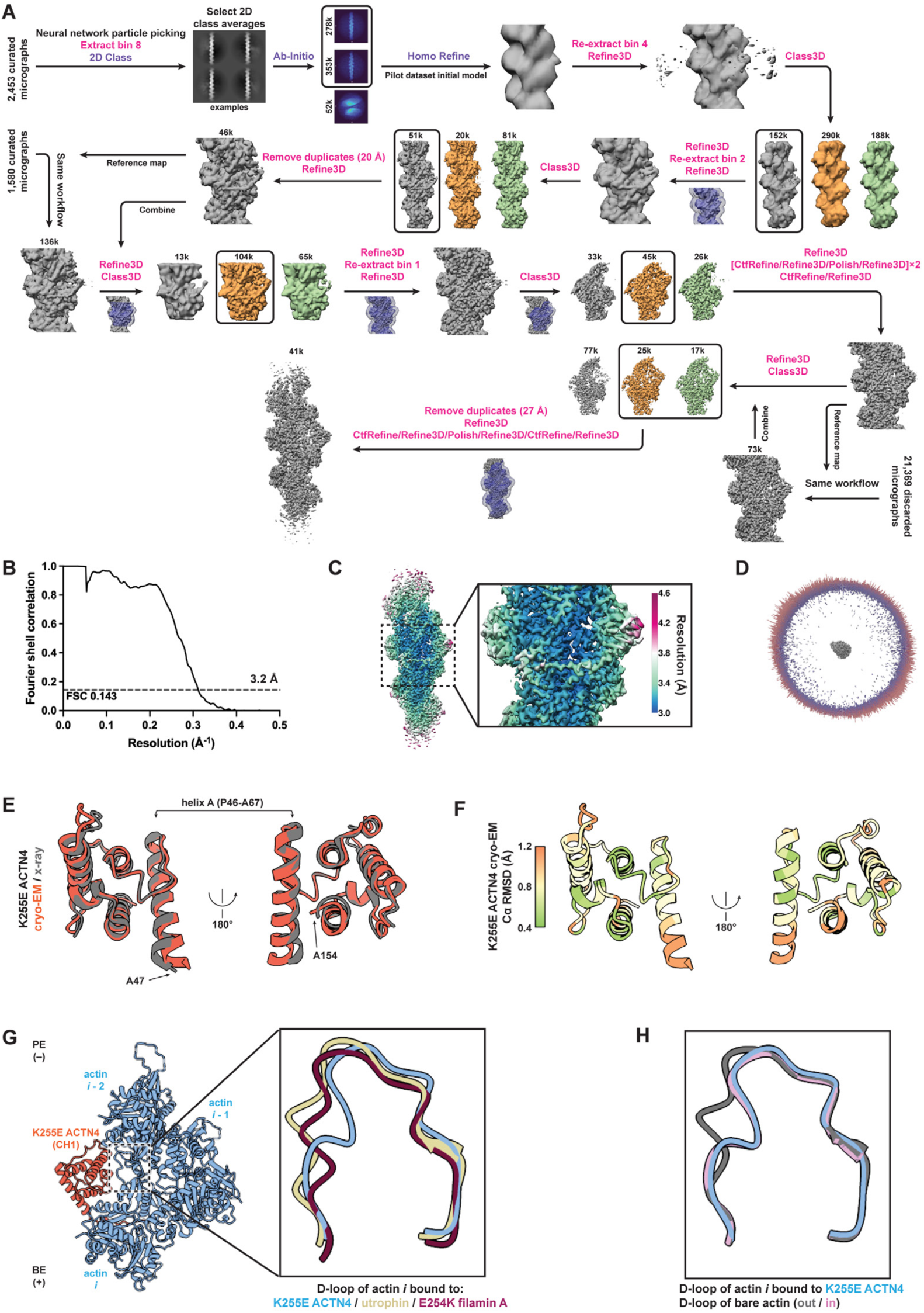
Cryo-EM data processing and analysis of K255E ACTN4–F-actin. **(A)** Data processing workflow. Each mask is displayed at the first step where it was used, superimposed on the corresponding reference. Jobs performed in RELION and cryoSPARC are indicated in magenta and purple text, respectively. **(B)** Fourier shell correlation (FSC) curve. **(C)** Local resolution assessment. **(D)** 3D angular distribution. **(E)** Superposition of K255E ACTN4 CH1 from the unbound ABD crystal structure^34^ (PDB 2R0O) and F-actin bound cryo-EM reconstruction (this study). **(F)** C_α_ root mean square deviation (RMSD) between superimposed structures from **E**.

**Figure S4:**
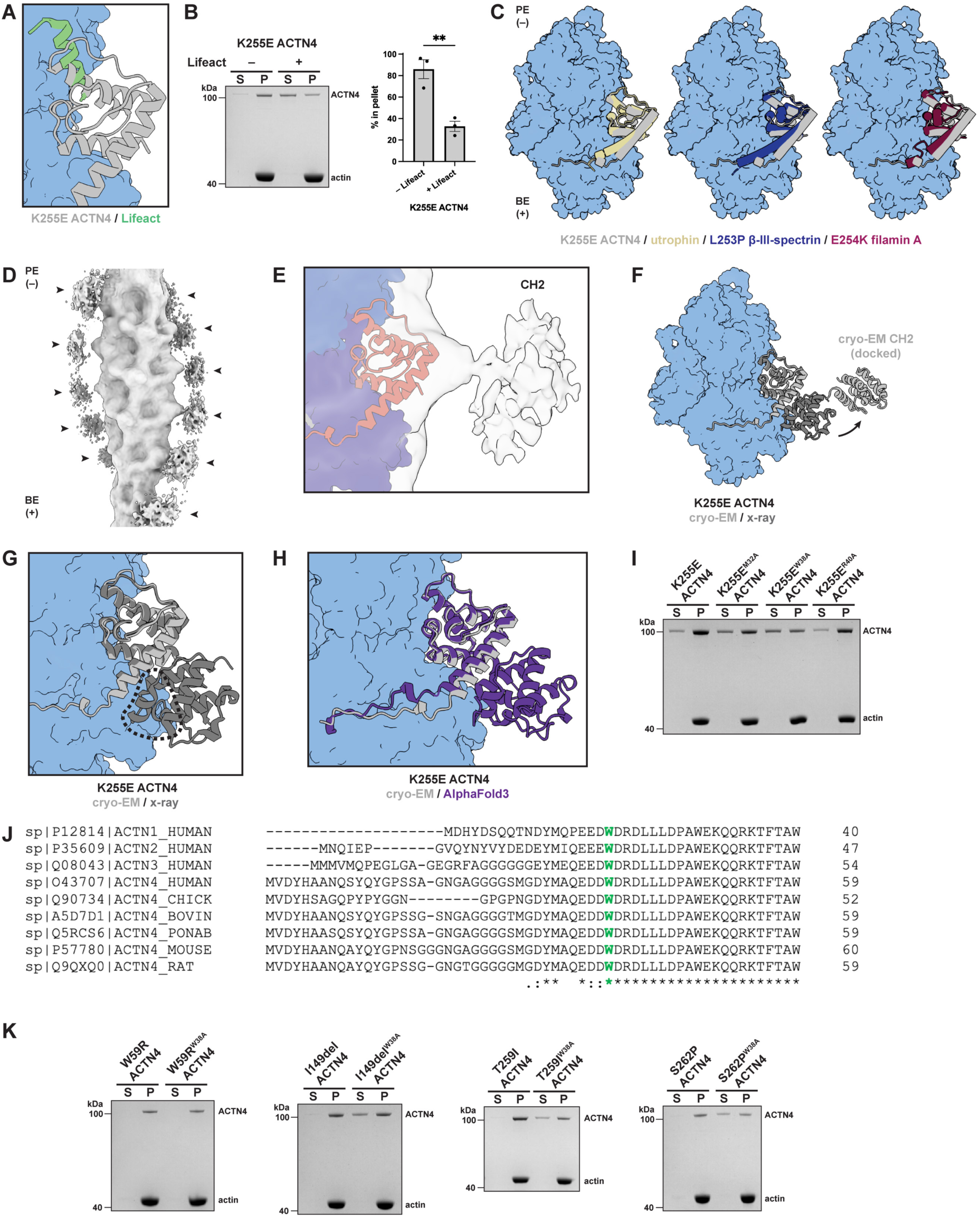
Additional analysis of the K255E ACTN4–F-actin structure. **(A)** Superposition of the Lifeact–F-actin complex^42^ (PDB 7AD9) and the K255E ACTN4–F-actin structure (this study) highlights their overlapping binding sites. **(B)** Representative SDS-PAGE and quantification of competition F-actin co-sedimentation assay between Lifeact and K255E ACTN4. S, supernatant. P, pellet. Upper band (105 kDa) is ACTN4, and lower band (42 kDa) is actin. Data are presented as mean ± SEM (*n* = 3). Conditions were compared by unpaired t test, \*\**P* < 0.01. **(C)** F-actin-bound utrophin^39^ (PDB 6M5G), L253P β-III-spectrin^37^ (PDB 6ANU), and E254K FLNA^36^ (PDB 6D8C) superimposed on F-actin-bound K255E ACTN4 (aligned on actin). Actins from the K255E ACTN4–F-actin structure are displayed. BE (+), barbed (plus) end; PE (−), pointed (minus) end. **(D)** Cryo-EM map derived from pilot K255E ACTN4–F-actin dataset, which featured excess K255E ACTN4 decoration. Diffuse densities consistent with displaced CH2 are highlighted (arrowheads). **(E)** K255E ACTN4–F-actin atomic model docked into pilot cryo-EM map from **D**, with unoccupied density attributable to CH2. **(F)** Approximate docking of CH2 into cryo-EM density alongside the K255E ACTN4–F-actin atomic model (light grey) highlights ABD opening relative to the unbound closed conformation^34^ (PDB 2R0O, superimposed on CH1). **(G)** Superposition of the unbound K255E ACTN4 ABD on F-actin-bound K255E ACTN4 (aligned on CH1) highlights region with clashes between CH2 and F-actin in the closed conformation. **(H)** AlphaFold3 predicted structure of the K255E ACTN4 ABD (residues 1-269) bound to F-actin (five human α-actin-1 subunits) superimposed the cryo-EM structure (aligned on actin). CH2 domain of the AlphaFold3 predicted structure remains in a closed conformation. The AlphaFold3-predicted NTE binding mode also differs from that in our cryo-EM structure. Residues 1-30 of K255E ACTN4 were disordered and extended away from F-actin in the predicted structure, and they are hidden for clarity. **(I)** SDS-PAGE of representative F-actin co-sedimentation assay for alanine mutations in the K255E ACTN4 NTE. Related to Fig. 1H. **(J)** Sequence alignment of NTEs of ACTN4 proteins from indicated species and human ACTN1-3. W38 (human ACTN4 numbering) is conserved across ACTNs (green). (**K**) SDS-PAGE of representative F-actin co-sedimentation assay for indicated FSGS-causing ACTN4 mutations and their corresponding W38A variants. Related to Fig. 1J.

**Figure S5:**
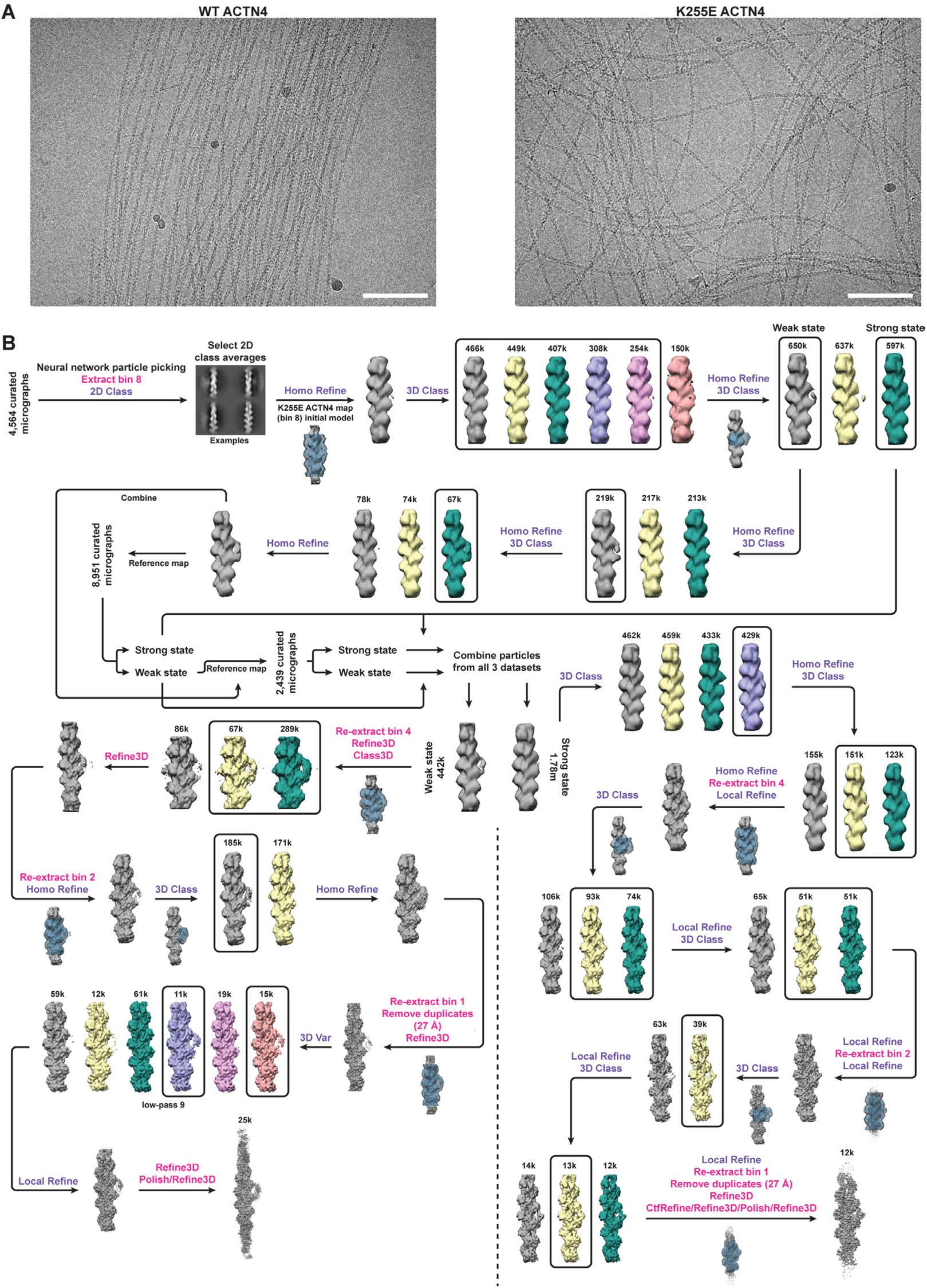
Cryo-EM and image processing for wild-type ACTN4–F-actin in the absence of myosin motors. **(A)** Representative micrographs from wild-type ACTN4–F-actin and K255E ACTN4–F-actin datasets, highlighting differential filament network organization. Scale bars, 100 nm. **(B)** Image processing workflow for recovering both weak and strong state cryo-EM maps. Each mask is displayed at the first step where it was used, superimposed on the corresponding reference. Jobs performed in RELION and cryoSPARC are indicated in magenta and purple text, respectively.

**Figure S6:**
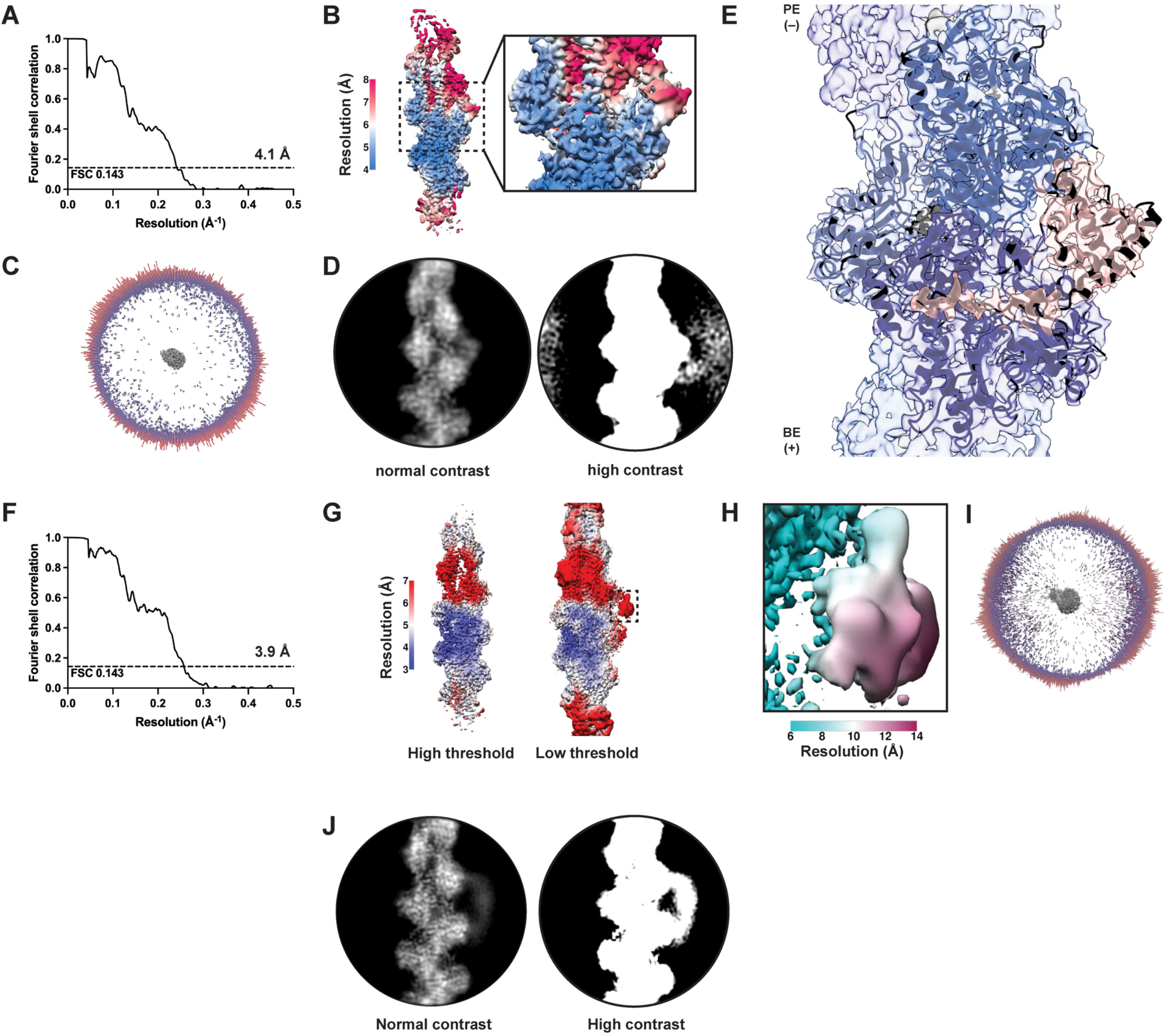
Cryo-EM data analysis for wild-type ACTN4–F-actin in the absence of myosin motors. **(A)** FSC curve for strong state map. **(B)** Local resolution assessment for strong state map. **(C)** 3D angular distribution of particles contributing to the strong state map. **(D)** 2D class average of particles contributing to the strong state map, viewed at different thresholds. At low threshold, diffuse density is visible that likely corresponds to CH2 (similar as visible for K255E ACTN4, Fig. 1D). **(E)** Atomic model of K255E ACTN4 bound to F-actin docked into the wild-type ACTN4 strong state map. BE (+), barbed (plus) end; PE (−), pointed (minus) end. **(F)** FSC curve for weak state map. **(G)** Overall local resolution assessment for weak state map. Note that color scale does not capture resolution range of boxed ABD region. **(H)** Local resolution assessment for ABD density in box from **G**, with adjusted color scale to capture resolution gradient in this region. **(I)** 3D angular distribution of weak state map. **(J)** 2D class average of particles contributing to the weak state map, viewed at different thresholds. Diffuse density distal from the filament surface is not present at low threshold (as is observed in the strong state), consistent with CH2 engaging F-actin in the weak state.

**Figure S7:**
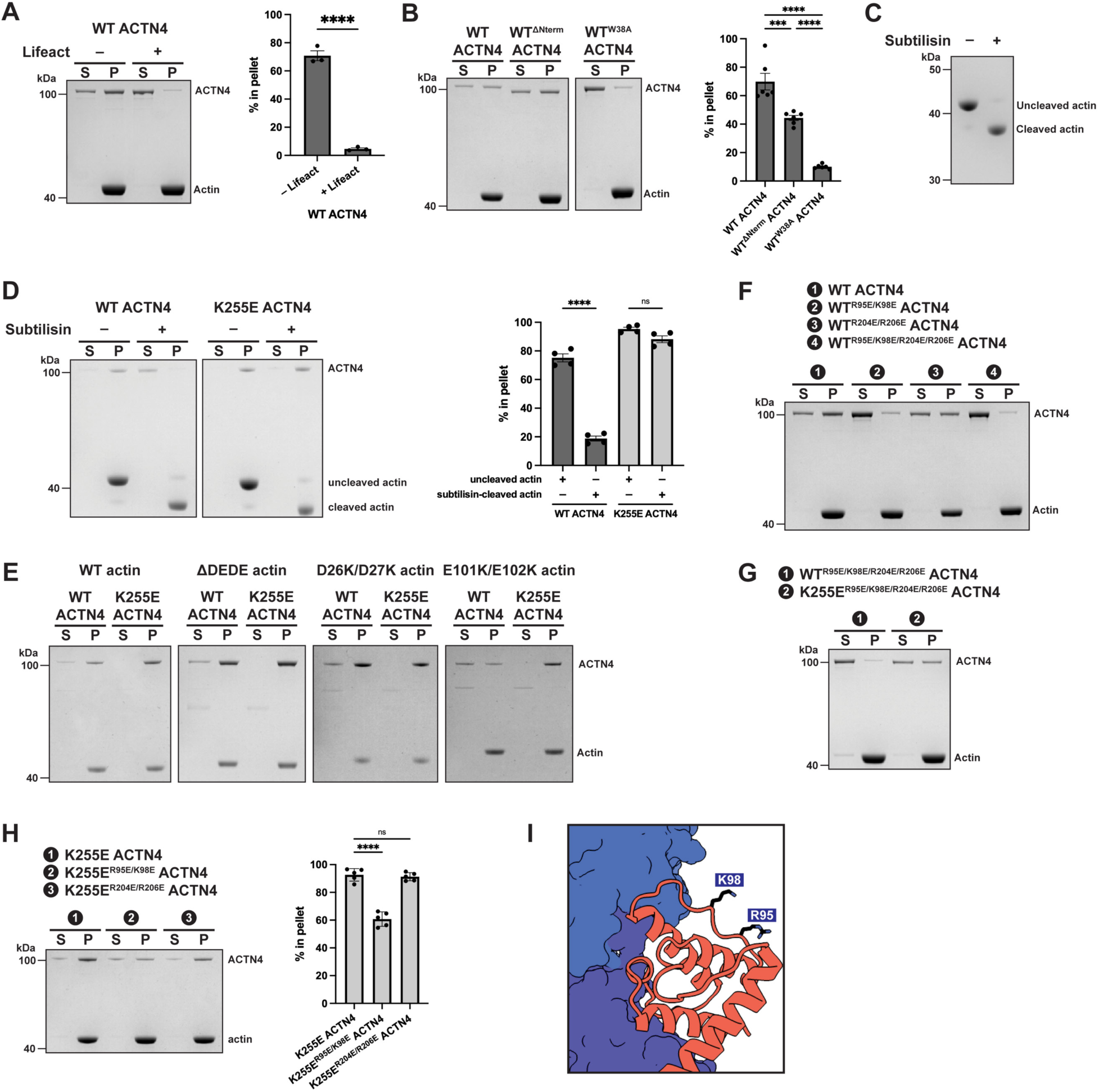
Biochemical assays to dissect F-actin binding modes of wild-type ACTN4. **(A)** SDS-PAGE of representative co-sedimentation assay and quantification of Lifeact competitive F-actin binding with wild-type ACTN4. S, supernatant. P, pellet. Upper band (105 kDa) is ACTN4, and lower band (42 kDa) is actin. Conditions were compared by unpaired t test (*n* = 3). **(B)** SDS-PAGE of representative co-sedimentation assay and quantification of F-actin binding by NTE point mutations and truncation in the context of wild-type ACTN4. Conditions were compared by ordinary one-way ANOVA with Tukey’s correction (*n* = 6). **(C)** SDS-PAGE of uncleaved and subtilisin-cleaved actin. **(D)** SDS-PAGE of representative co-sedimentation assay and quantification of uncleaved versus subtilisin-cleaved F-actin binding by wild-type and K255E ACTN4. Conditions were compared by ordinary two-way ANOVA with Šídák’s correction (*n* = 4). **(E)** Representative SDS-PAGE of wild-type and K255E ACTN4 co-sedimentation assay with F-actin featuring charge reversal mutations (in recombinant human α-actin-1). Related to Fig. 2D. **(F)** Representative SDS- PAGE of F-actin co-sedimentation assay for ACTN4 charge reversal mutations in the context of the wild-type ACTN4 background. Related to Fig. 2G. **(G)** Representative SDS-PAGE of F-actin co-sedimentation assay for simultaneous charge reversal mutations in both CHDs of wild-type and K255E ACTN4. Related to Fig. 2H. **(H)** Representative SDS-PAGE of F-actin co-sedimentation assay and quantification for charge reversal mutations in K255E ACTN4. Conditions were compared by ordinary one-way ANOVA with Dunnett’s correction (*n* = 5). **(I)** Residues contributing to the positive patch on CH1 are displayed on the K255E ACTN4–F-actin structure, highlighting how they are facing away from the F-actin surface in the strong state. For quantification of co-sedimentation assays throughout the figure, data are presented as mean ± SEM. \*\*\**P* < 0.001, \*\*\*\**P* < 0.0001, ns = not significant (*P* ≥ 0.05)

**Figure S8:**
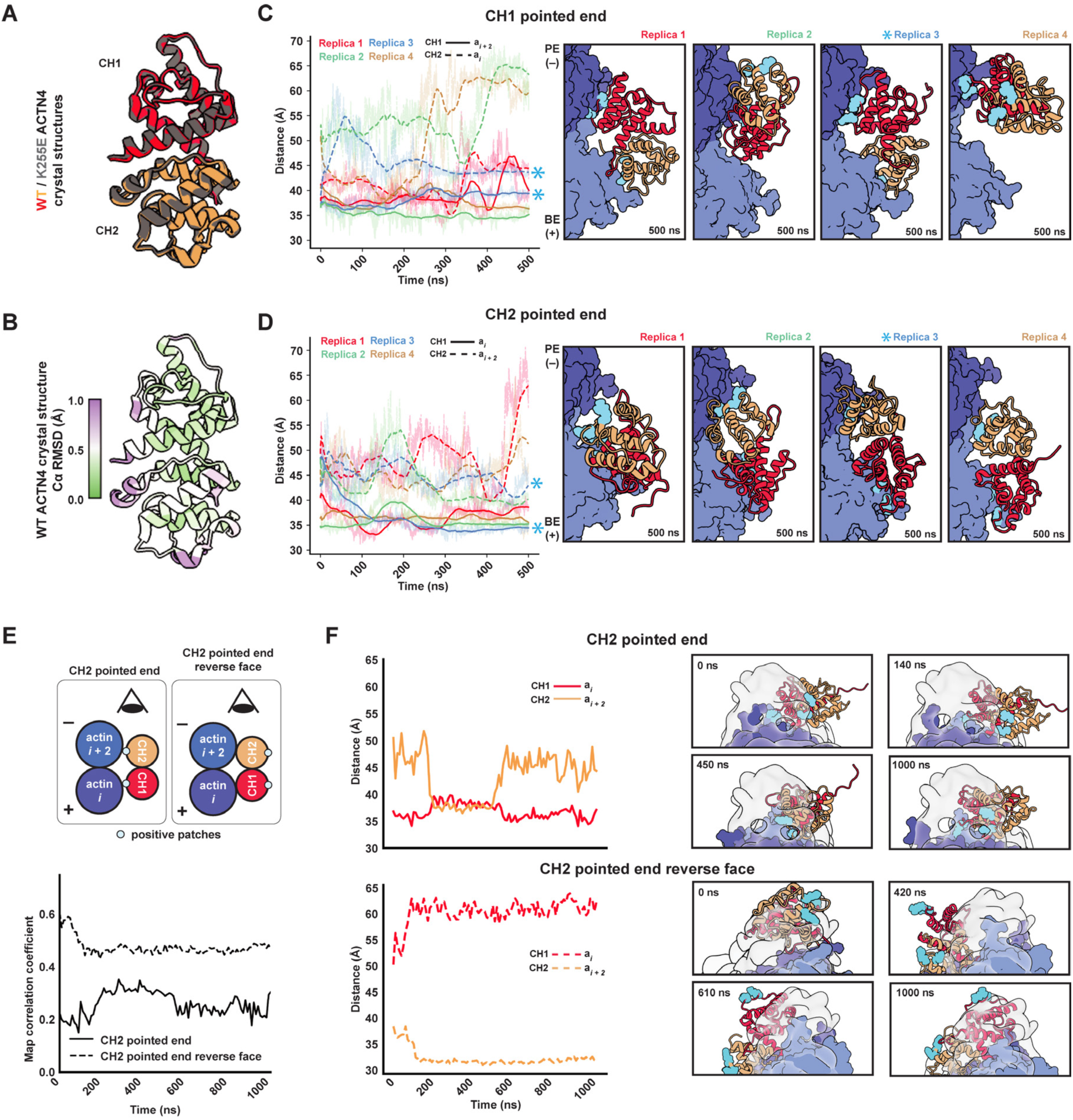
Additional MD analysis of the weak-binding state. **(A)** Superposition of wild-type^35^ (PDB 6O31) and K255E^34^ (PDB 2R0O) ACTN4 ABD crystal structures. **(B)** C_α_ RMSD between superimposed structures from **A**. **(C-D)** Plots of CH domain distance to the nearest actin across four replica simulations (left) and snapshots corresponding to the final frame of each 500 ns trajectory (right) for the CH1 pointed end **(C)** and CH2 pointed end **(D)** orientations. Replicas indicated with asterisks were run for an additional 1 µs. BE (+), barbed (plus) end; PE (−), pointed (minus) end. **(E)** Cartoon of additional ACTN4 ABD orientations examined in MD simulations of weak state (top). Map correlation coefficients from 1 µs MD simulations of ACTN4 ABD in orientations from (bottom). (**F**) Plots of CH domain distance to the nearest actin subunit (left) and representative simulation snapshots overlaid with the cryo-EM map (right) of the CH2 pointed end (top) and CH2 pointed end reverse face (bottom) orientations. Snapshots are from the view indicated in **E**. Distances were calculated between the centers of mass of C_α_ atoms. Cyan residues indicate positive amino acids highlighted in Fig. 2B.

**Figure S9:**
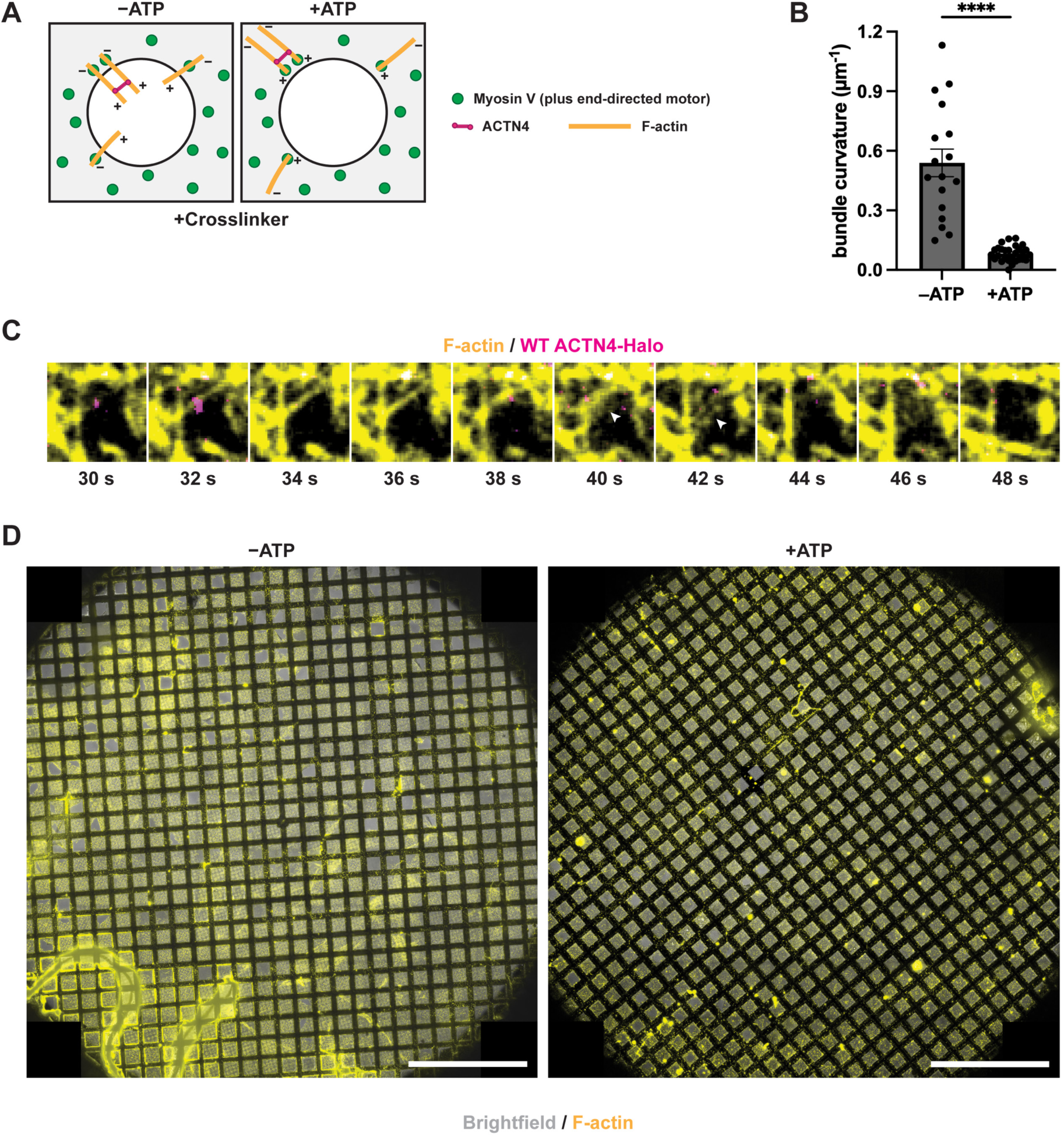
Force reconstitution assay on cryo-EM grids. **(A)** Schematic of force reconstitution assay behavior when crosslinkers form parallel F-actin bundles, which are anticipated to glide on to the carbon film in a manner similar to unbundled filaments. **(B)** Example of force-induced bundle ruptures (arrowheads), which occurred at later time points in the hole displayed in Fig. 3C (bottom). **(C)** Quantification of bundle curvature from epifluorescence of force reconstitution assays in the presence of wild-type ACTN4-Halo with or without ATP. Analysis was performed on 5 and 2 movies for −ATP and +ATP conditions, respectively. Data are presented as mean ± SEM (*n* = 17 and 28 for −ATP and +ATP conditions, respectively). Conditions were compared by unpaired t test, \*\*\*\**P* < 0.0001. **(D)** Cryo-fluorescence images of −ATP and +ATP grids. Whole-grid atlases were generated by maximum intensity projection and stitching of images of smaller regions. Brightfield and F-actin (Alexa Fluor Plus 555 phalloidin) channels are displayed. Scale bars, 500 µm.

**Figure S10:**
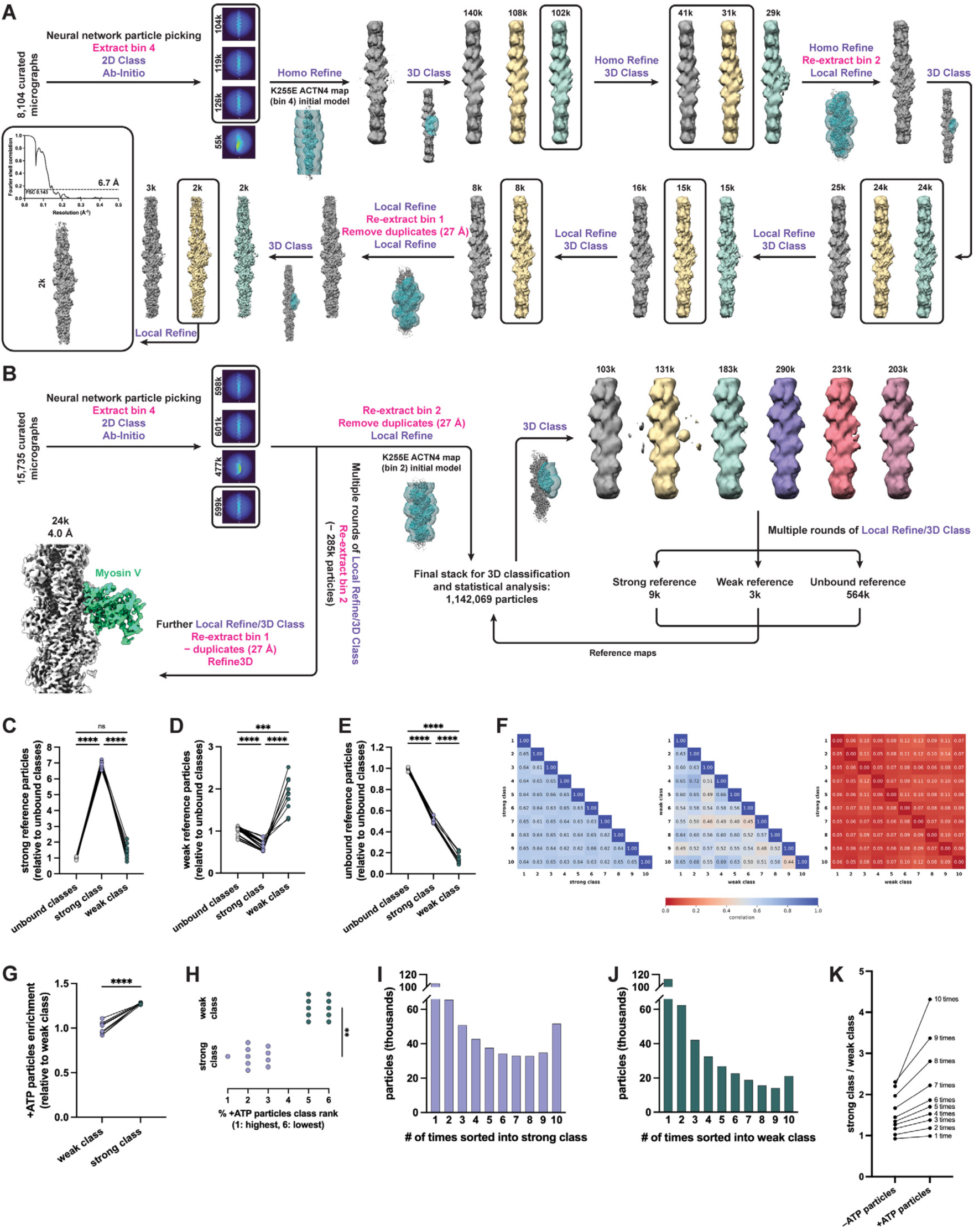
Cryo-EM data processing and analysis for wild-type ACTN4–F-actin in the presence of myosin motor forces. **(A)** Image processing workflow for analyzing wild-type ACTN4 bound to F-actin from the +ATP dataset alone. Each mask is shown at the first step it was used, superimposed on the corresponding reference. The FSC curve for the final reconstruction is also displayed. **(B)** Detailed workflow for processing combined −ATP and +ATP datasets in preparation for quantification by 3D classification, featuring data cleaning and generation of references for strong, weak, and unbound states. Jobs performed in RELION and cryoSPARC are displayed in magenta and purple, respectively. **(C-E)** Post-supervised 3D classification assignments of particles originally associated with strong (**C**), weak (**D**), and unbound (**E**) references. Data are normalized to class sizes and expressed as fold-enrichment relative to unbound classes. Conditions were compared by repeated measures one-way ANOVA with Geisser-Greenhouse and Tukey’s corrections. **(F)** Correlation matrices assessing similarity of class assignments between independent 3D classification runs. Analysis is based on number of shared particles, normalized to class size displayed on horizontal axis. **(G)** Quantification of +ATP dataset particle enrichment in weak and strong classes (expressed as fold change relative to weak class). Conditions were compared by ratio paired t test. **(H)** For each 3D classification run, individual classes were ranked by percent of particles derived from the +ATP dataset and plotted. Rankings for strong and weak classes within runs were then compared by Wilcoxon matched-pairs signed rank test. **(I-J)** Quantification of per-particle consistency of class assignment across classification runs for the strong **(I)** and weak **(J)** classes. **(K)** Analysis of the relationship between how consistently particles appear in their respective class (I and J) and the shift in class assignments between -ATP and +ATP datasets. A larger magnitude ATP induced shift towards the strong class is observed for more consistently assigned particles. Related to Fig. 3G. For all statistical comparisons throughout the figure, *n* = 10 independent 3D classification runs. \*\*\*\**P* < 0.0001, ns = not significant (*P* ≥ 0.05).

**Figure S11:**
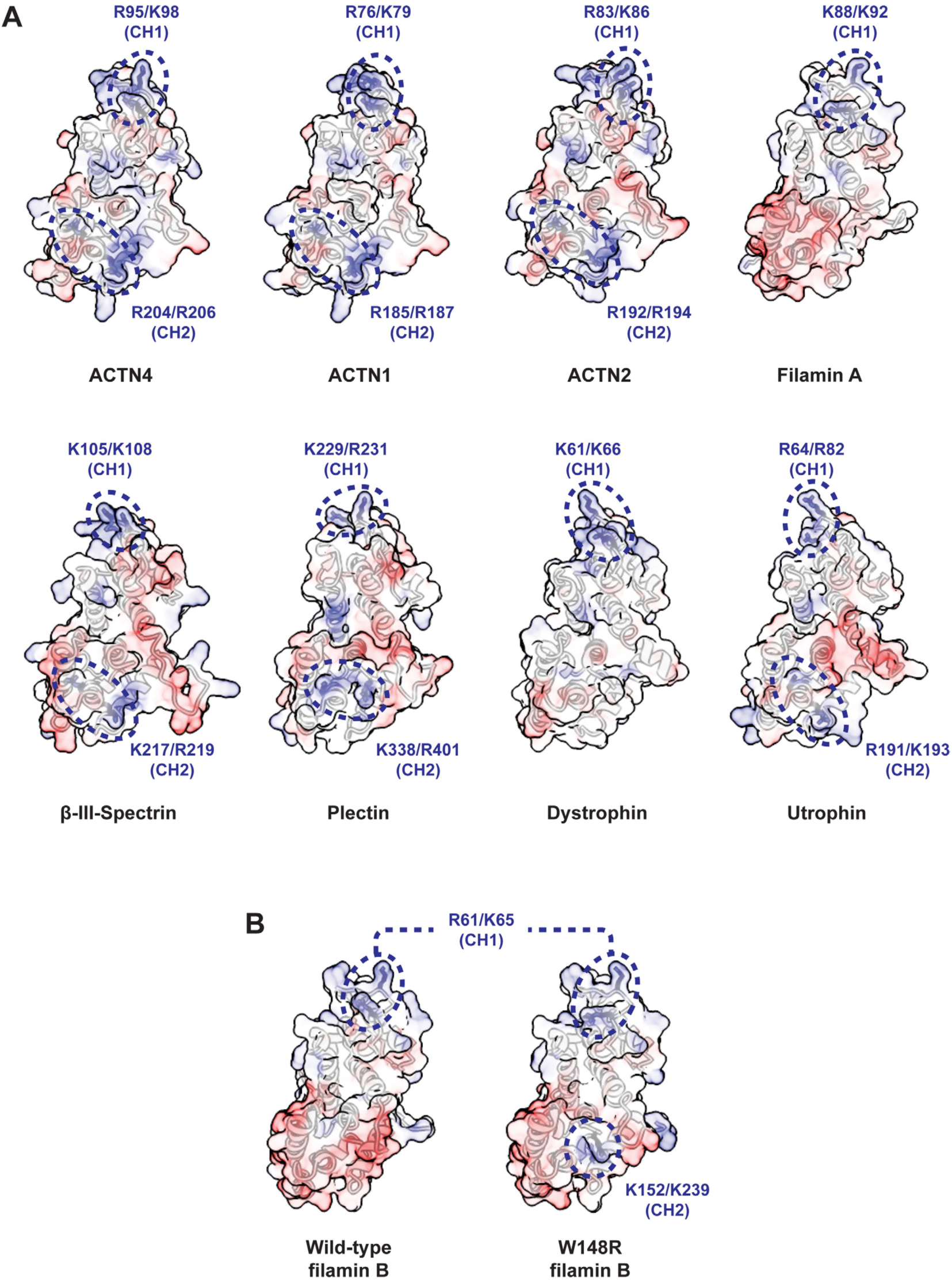
Charge distributions across tandem CHD ABDs. **(A)** Comparison of electrostatic potential surface maps of wild-type ACTN4^35^ (PDB 6O31), ACTN1^32^ (PDB 2EYI), ACTN2^90^ (PDB 4D1E), filamin A^33^ (PDB 3HOP), β-III-spectrin^91^ (AlphaFold DB/Uniprot O15020), plectin^92^ (PDB 1MB8), dystrophin^93^ (PDB 9D58), and utrophin (AlphaFold3 prediction of residues 31-255) ABDs. **(B)** Comparison of electrostatic potential surface maps of wild-type (PDB 2WA5) and W148R (PDB 2WA6) filamin B ABDs^62^.

## Movie Legends

**Movie S1: MD simulations of the weak state.** CH1 pointed end and its reverse face simulations are those analyzed in Fig. 2I-J. CH2 pointed end and its reverse face simulations are those analyzed in Fig. S8E-F. For each condition, 1 µs of MD simulation is displayed, with snapshots every 10 ns.

**Movie S2: Epifluorescence of force reconstitution assay in the absence of ACTN4, without and with ATP.** Related to Fig. 3B.

**Movie S3: Epifluorescence of force reconstitution assay in the presence of ACTN4, without and with ATP.** Related to Fig. 3C.

---

**Table S1:** Cryo-EM data collection, refinement, and validation statistics.

|  | K255E ACTN4<br>bound to F-actin<br>EMDB: XXXX<br>PDB: XXXX | Wild-type ACTN4<br>bound to F-actin<br>(strong state)<br>EMDB: XXXX | Wild-type ACTN4<br>bound to F-actin<br>(weak state)<br>EMDB: XXXX | Wild-type ACTN4<br>bound to F-actin<br>under force (+ATP<br>dataset only)<br>EMDB: XXXX |
| --- | --- | --- | --- | --- |
| **Data collection and processing** |  |  |  |  |
| Microscope | Titan Krios | Titan Krios | Titan Krios | Titan Krios |
| Detector | K3 | K3 | K3 | K3 |
| Magnification | 81,000 | 81,000 | 81,000 | 81,000 |
| Voltage (kV) | 300 | 300 | 300 | 300 |
| Electron exposure (e<sup>-</sup>/Å<sup>2</sup>) | 49.238 | 49.238 | 49.238 | 49.238 |
| Defocus range (μm) | -0.8 to -2.2 | -0.8 to -2.2 | -0.8 to -2.2 | -0.8 to -2.2 |
| Pixel size (Å) | 1.09 | 1.09 | 1.09 | 1.09 |
| Symmetry imposed | C1 | C1 | C1 | C1 |
| Micrographs (no.) | 25,402 | 15,954 | 15,954 | 8,104 |
| Initial particle images (no.) | 7,291,435 | 9,389,999 | 9,389,999 | 1,734,836 |
| Final particle images (no.) | 41,450 | 25,362 | 12,309 | 2,078 |
| Map resolution (Å) | 3.2 | 4.1 | 3.9 | 6.7 |
| FSC threshold | 0.143 | 0.143 | 0.143 | 0.143 |
| Map resolution range (Å) | 3.0–5.8 | 4.2–27.1 | 3.5–18.0 | 4.2–16.2 |
| **Refinement** |  |  |  |  |
| Initial model used<br>(PDB code) | 8D14, 2R0O | - | - | - |
| Model resolution (Å) | 3.2 | - | - | - |
| FSC threshold | 0.5 | - | - | - |
| Map sharpening<br>*B* factor (Å<sup>2</sup>) | -41.7 | -49.5 | -50.6 | - |
| Model composition | 3 actin<br>protomers,<br>1 K255E ACTN4 | - | - | - |
| Non-hydrogen atoms | 9855 | - | - | - |
| Protein residues | 1243 | - | - | - |
| Ligands | 3 Mg<sup>2+</sup>, 3 ADP,<br>3 PO<sub>4</sub><sup>3-</sup>, 1<br>phalloidin | - | - | - |
| ***B* factors (Å<sup>2</sup>)** |  |  |  |  |
| Protein | 40.57 | - | - | - |
| Ligand | 27.24 | - | - | - |
| **RMSD** |  |  |  |  |
| Bond lengths (Å) | 0.002 | - | - | - |
| Bond angles (°) | 0.545 | - | - | - |
| **Validation** |  |  |  |  |
| MolProbity score | 1.21 | - | - | - |
| Clashscore | 3.78 | - | - | - |
| Poor rotamers (%) | 0.67 | - | - | - |
| **Ramachandran plot** |  |  |  |  |
| Favored (%) | 97.79 | - | - | - |
| Allowed (%) | 2.21 | - | - | - |
| Disallowed (%) | 0.00 | - | - | - |

